# Deep-body feelings: ingestible pills reveal gastric correlates of emotions

**DOI:** 10.1101/2023.02.17.528509

**Authors:** Giuseppina Porciello, Alessandro Monti, Maria Serena Panasiti, Salvatore Maria Aglioti

## Abstract

Although it is generally held that gastro-intestinal (GI) signals are related to emotions, direct evidence for such a link is currently lacking. One of the reasons why the internal milieu of the GI system is poorly investigated is because visceral organs are difficult to access and monitor. To directly measure the influence of endoluminal markers of GI activity on the emotional experience, we asked a group of healthy male participants to ingest a pill that measured pH, pressure, and temperature of their GI tract while they watched video-clips that consistently induced disgust, fear, happiness, sadness, or a control neutral state. In addition to the objective physiological markers of GI activity, subjective ratings of perceived emotions and visceral (i.e. gastric, respiratory and cardiac) sensations were recorded. We found that when participants observed fearful and disgusting video-clips, they reported to perceive not only cardiac and respiratory sensations but also gastric sensations, such as nausea. Moreover, we found that there was a clear relation between the physiology of the stomach and the perceived emotions. Specifically, when disgusting video-clips were displayed, the more acidic the pH, the more participants reported feelings of disgust and fear; the less acidic the pH, the more they reported happiness. Our findings suggest that gastric signals contribute to unique emotional states and that ingestible pills may open new avenues for exploring the deep-body physiology of emotions.

## Introduction

Whether specific patterns of physiological signals coming from visceral organs trigger a unique emotional state is a hotly debated question (James, 1894). Somatic theories of emotions posit that afferent physiological signals are essential for experiencing distinct emotional states (Damasio, 1999; Harrison et al., 2010; James, 1894). However, so far the large majority of the studies on the influence of visceral signals on emotions focused on the role of cardiac and respiratory activity (Kreibig, 2010; Rainville et al., 2006), neglecting the potential role of the gastrointestinal (GI) tract. Yet, the GI system is directly connected with the central nervous system via vagal and spinal afferents (Azzalini et al., 2019). In turn, different cortical areas directly influence parasympathetic and sympathetic control of the stomach (Levinthal and Strick, 2020). Moreover, it has been shown that the slow electrical rhythm (∼0.05 Hz) generated in the stomach by the interstitial cells of Cajal interacts with resting-state neural networks (Rebollo et al., 2018). This interaction, described as the “gastric network” and mostly composed by sensory-motor regions (Rebollo and Tallon-Baudry, 2022), seems to be stronger during stress (Jeanne et al., 2022), and to go beyond food intake and weight regulatory functions (Levakov et al., 2021), possibly playing a role in higher-order cognitive and emotional functions (Azzalini et al., 2019; Porciello et al., 2018; Rebollo et al., 2021). That the GI system plays a fundamental role in the experience of emotions has been suggested by studies in which healthy participants were asked to associate basic and complex emotions with the subjective perception of changes taking place in specific parts of the body, including the heart, the lungs, and the GI tract (Nummenmaa et al., 2018, 2014). Tellingly, people tend to link changes of activity in the stomach and the bowel with the experience of disgust and happiness (Nummenmaa et al., 2014). It is worth noting that, although insightful, these findings derive from mere self-report data. Studies of objective indices of association between deep body activation and emotions are meager. To the best of our knowledge, these few studies show a direct contribution of the GI system to the emotional experience of disgust (Harrison et al., 2010; Shenhav and Mendes, 2014) using electrogastrography (EGG), a technique that allows to record the myoelectric activity of the stomach through electrodes applied on the skin overlying the epigastric region. Overall, these experiments suggest that the gastric myoelectric dysrhythmias contribute to the emotional experience of disgust (Harrison et al., 2010; Shenhav and Mendes, 2014) and that disgusting stimuli specifically reduce slow gastric activity (bradygastria) and increase normal (normogastria) and high (tachygastria) gastric activity (Gianaros et al., 2001; Peyrot Des Gachons et al., 2011; Shenhav and Mendes, 2014; Stern et al., 2001; Vianna and Tranel, 2006). In line with these findings, a recent randomized, placebo-controlled study (Nord et al., 2021) run on healthy participants showed that oculomotor avoidance (a marker of disgust) evoked by disgusting stimuli significantly decreases after taking domperidone, an antiemetic and prokinetic agent that seems to reduce gastric dysrhythmias. Unfortunately, though, no EGG or any other objective measure of gastric physiology was provided. One of the reasons why the endoluminal milieu of the GI system is poorly known is because internal organs are difficult to access and monitor using non-invasive or minimally invasive tools. To overcome this limitation, recent technologies, such as ingestible sensors, have been developed in animal (e.g. Mimee et al., 2018) and clinical research for non-invasive diagnostics of diseases affecting the GI system (Mandsberg et al., 2020; Min et al., 2020; Rao et al., 2011) and for local drug delivery (Mandsberg et al., 2020). Interestingly, recent findings show that ingestible pills can be used to explore higher-order cognitive and emotional processes (Mayeli et al., 2021; Monti et al., 2022). In particular, we have provided the first evidence (Monti et al., 2022) that specific patterns of GI signals covary with specific facets of bodily self-consciousness (feelings of body location, agency, and disembodiment). In the present study, we used inert ingestible pills (SmartPills) that gauge pH, core temperature and pressure in different segments of the GI tract (Saad, 2016) while participants watched a validated set of short video-clips that reliably induced four basic emotions: fear, disgust, sadness, and happiness (Tettamanti et al., 2012), see **Fig.1** for a graphical illustration of the emotional induction procedure. The gastric internal parameters detected through SmartPills were complemented with standard surface electrogastrography (Yin and Chen, 2013). Our approach allowed us to identify internally and externally recorded gastric markers associated with specific emotional experiences. We also collected self-report ratings of emotions, visceral feelings, and arousal. In this way, we checked the efficacy of our stimuli and measured how the emotional experience induced by the video-clips made our participants aware of their visceral state (Craig, 2003). As far as self-report ratings are concerned, we expected that: i) in accordance with previous findings (Tettamanti et al., 2012) all the emotional video-clips would trigger the intended emotion; ii) basic emotions would be associated with specific visceral sensations mirroring results coming from the cardiac and respiratory domains (e.g. Rainville et al., 2006); iii) participants would report higher gastric sensations after observing disgusting and happy video-clips as suggested by Nummenmaa and collaborators (Nummenmaa et al., 2014); and iv) there would be no differences in self-reported arousal ratings between the emotional video-clips, in line with previous literature (e.g. Posner et al., 2005; Russell and Feldman Barrett, 1999). With respect to the GI physiological signals, our hypotheses relied on a limited number of prior studies that did not make use of the pill technology. Hence, the present study should be considered as the first attempt to explore the neglected contribution of GI internal markers to the emotional experience. On the basis on the published literature on core disgust (Harrison et al., 2010; Nord et al., 2021; Shenhav and Mendes, 2014), we hypothesized that the subjective experience of disgust during the observation of the disgusting video-clips would be linked to individual gastric rhythm as indexed by the EGG traces and the GI pressure recorded by the ingestible pill. Moreover, based on single case studies (Bennett and Venables, 1920; Wolf and Wolff, 1947), we also predicted that the subjective experience of disgust, fear, and sadness would be associated with more acidic pH in the stomach. Finally, based on the literature showing that externally recorded oral or axillary body temperature increases after the observation of disgusting and stressful stimuli (Marazziti et al., 1992; Stevenson et al., 2012), we predicted that, compared with happy and neutral emotional states, disgust, fear, and sadness,s triggered by the corresponding video-clips, would be associated with an increase of GI temperature.

**Fig.1.**
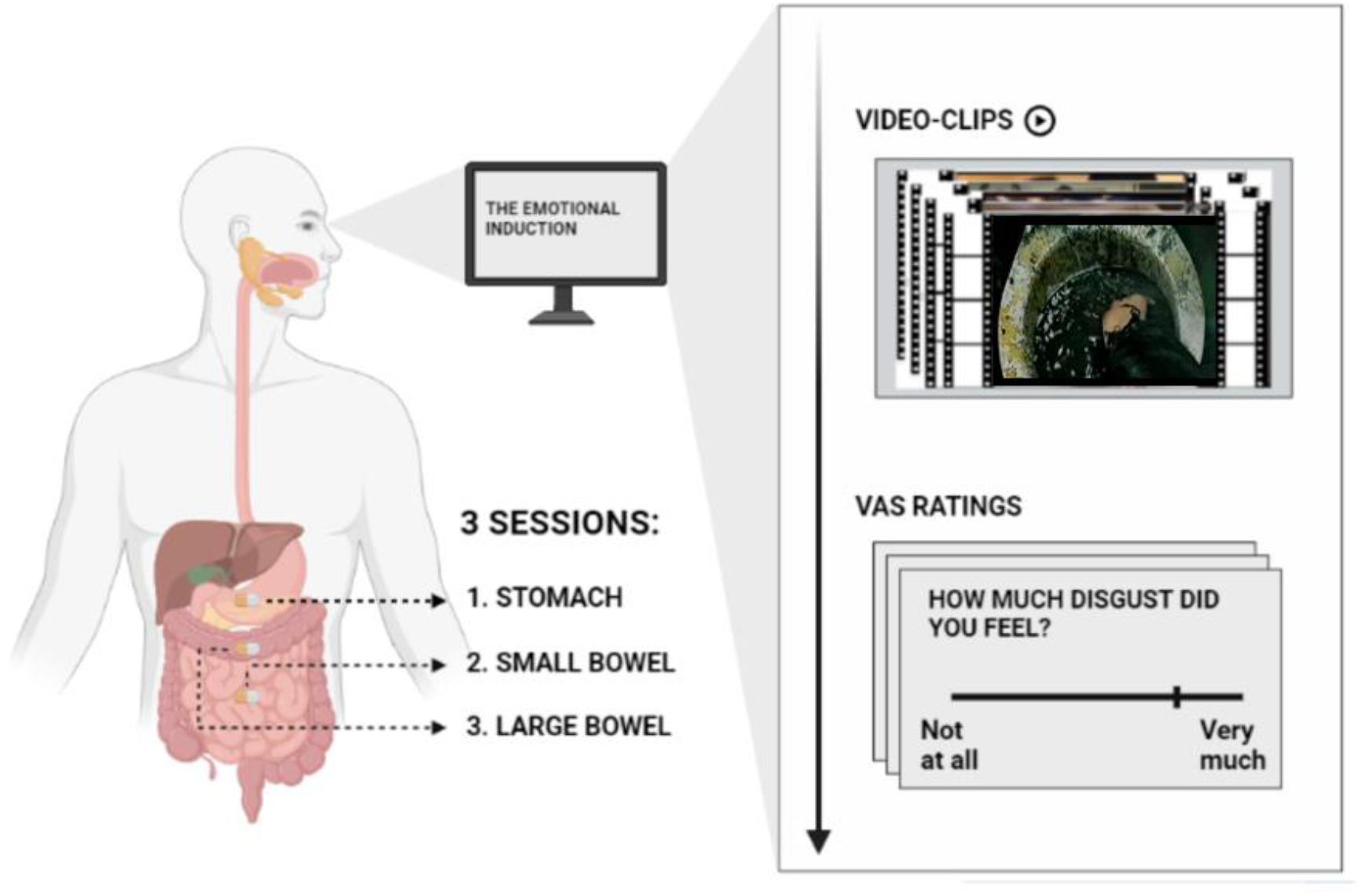
Illustration of the emotional induction procedure. Participants were asked to observe five blocks of 24 video-clips each. Four blocks were associated with a specific emotion, namely, happiness, disgust, sadness, and fear. An additional block consisting of neutral video-clips served as control. Video-clips were edited color soundless film excerpts lasting ∼9 s each and were selected and validated by Tettamanti and collaborators (2012). After each block, participants were asked to respond to an 8-items questionnaire by positioning a computer mouse on 0 (“Not at all”) – 100 (“Very much”) visuo-analogue scales (VAS). The emotional induction procedure was repeated three times, each corresponding to the ingestible pill position in the stomach (1) in the small (2) and in the large bowel (3). This figure was designed with BioRender.com and edited with Adobe Photoshop® 7.0.

## Results

### Subjective emotional experience

#### Capsule in the stomach (Session 1)

The Friedman ANOVA comparing the perceived emotional experience (i.e. disgust, fear, happiness and sadness) triggered by the five types of video-clips (varying for their content, i.e. disgusting; fearful; happy; sad; and neutral) in the first session (namely when the capsule was in the stomach) was statistically significant (χ^2^ (19) = 296.91; p < 0.0001), suggesting that participants perceived different emotions after observing the different content of the video-clips. Specifically, planned post-hoc Bonferroni-corrected Wilcoxon matched-pairs tests showed that disgust was primarily perceived after watching disgusting video-clips, i.e. the VAS ratings of perceived disgust given after observing disgusting video-clips were higher than those given after observation of all the other video-clips (all *Z*s ≥ 4.372; all *p*s ≤ 0.0001). Fear was primarily perceived after watching fearful video-clips, i.e. the VAS ratings of perceived fear given after fearful video-clips were higher than after the other video-clips (all *Z*s ≥ 2.972; all *p*s ≤ 0.003). Happiness was primarily perceived after watching happy video-clips, i.e. the VAS ratings of perceived happiness given after happy video-clips were higher than after the other video-clips (all *Z*s ≥ 4.445; all *p*s ≤ 0.0001). Finally, sadness was primarily perceived after watching sad video-clips, i.e. the VAS ratings of perceived sadness given after happy video-clips were higher than the other video-clips (all Zs ≥ 4.422; all *p*s ≤ 0.0001). All the significant post-hoc comparisons are shown in **Fig.2**.

**Fig.2.**
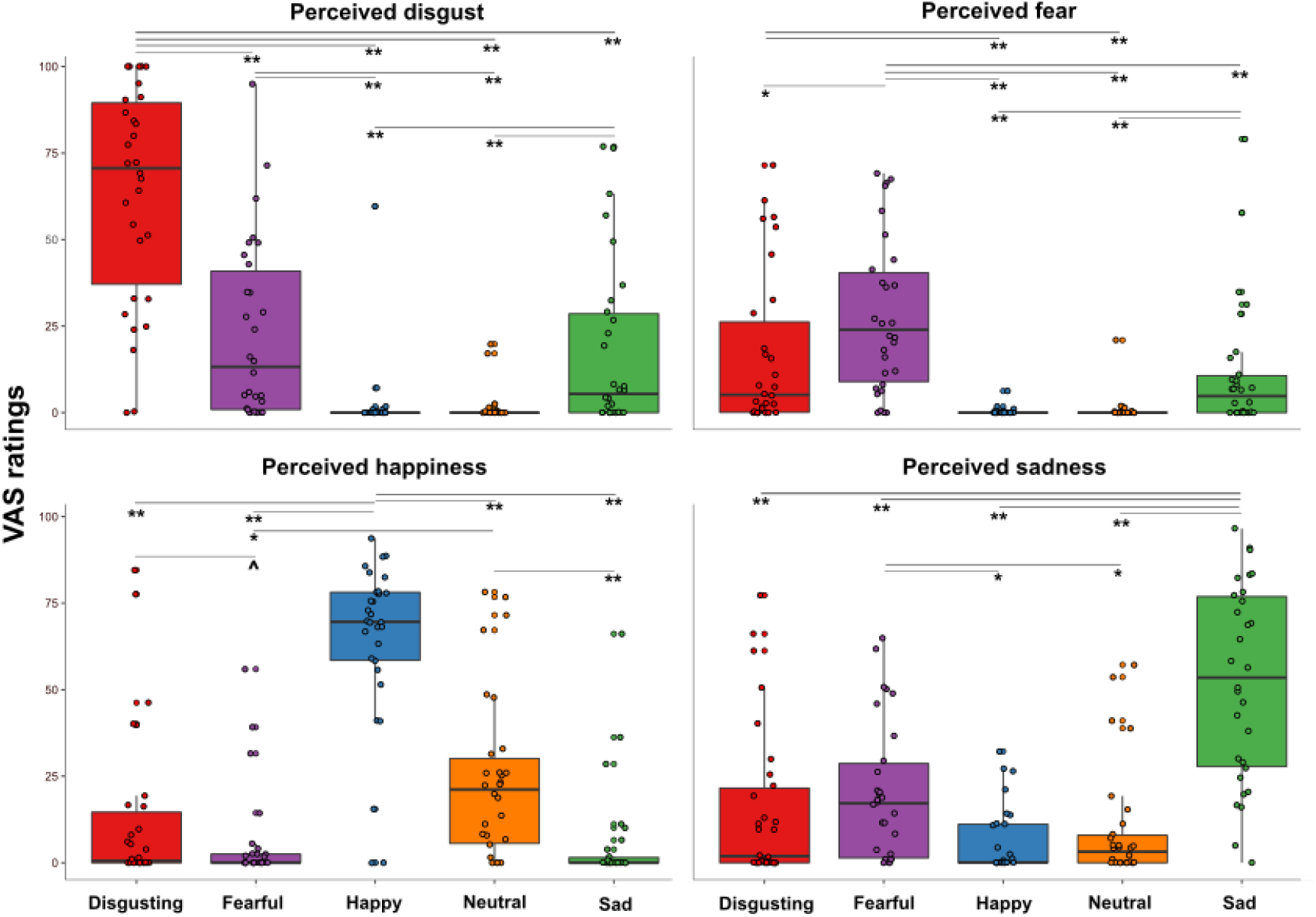
Perceived emotions results (pill in the stomach, session 1). Perceived emotions (disgust, fear, happiness, and sadness) measured using 0-100 visuo-analogue scale (VAS) ratings, as a function of the five categories of video-clips (disgusting, fearful, happy, sad, and neutral) shown during the first session of this study (i.e., when the capsule was in the stomach). Significant differences refer to Bonferroni-corrected, Wilcoxon matched-pairs tests (threshold for multiple comparisons was set at 0.0125). ∧p = 0.013; * p < 0.01; ** p ≤ 0.001.

#### Capsule in the small bowel (Session 2)

The Friedman ANOVA comparing the perceived emotional experience triggered by the five types of video-clips in the second session (namely when the capsule was in the small bowel) was statistically significant (χ^2^(19) = 358.763; *p* ≤ 0.0001), suggesting that participants perceived different emotions after observing the different content of the video-clips. Planned post-hoc Bonferroni-corrected Wilcoxon matched-pairs tests showed results similar to those found in the first session. Specifically, VAS ratings of perceived disgust given after disgusting video-clips were higher than those given after all the other video-clips (all *Z*s ≥ 4.597; all *p*s ≤ 0.0001). VAS ratings of perceived fear given after fearful video-clips were higher than those given after all the other video-clips (all *Z*s ≥ 3.029; all *p*s ≤ 0.002). VAS ratings of perceived happiness given after happy video-clips were higher than those given after all the other video-clips (all *Z*s ≥ 4.372; all *p*s ≤ 0.0001). Finally, VAS ratings of perceived sadness given after sad video-clips were higher than those given after all the other video-clips (all Zs ≥ 4.509; all *p*s ≤ 0.0001). Due to the fact that session 2 results mirror those of session 1, they are plotted in the Supplementary Information (**Fig.S1**).

#### Capsule in the large bowel (Session 3)

The Friedman ANOVA comparing the perceived emotional experience triggered by the five types of video-clips when the capsule was in the large bowel was statistically significant (χ^2^(19) = 328.053; *p* ≤ 0.0001), suggesting that participants perceived different emotions even after observing for three times the different content of the same video-clips. Planned post-hoc Bonferroni-corrected Wilcoxon matched-pairs tests showed results akin to those found in the first and second session. Specifically, VAS ratings of perceived disgust given after disgusting video-clips were higher than those given after all the other video-clips (all *Z*s ≥ 4.421; all *p*s ≤ 0.0001). VAS ratings of perceived fear given after fearful video-clips were higher than those given after all the other video-clips (all *Z*s ≥ 3.484; all *p*s ≤ 0.0005). VAS ratings of perceived happiness given after happy video-clips were higher than those given after all the other video-clips (all *Z*s ≥ 4.379; all *p*s ≤ 0.0001). Finally, VAS ratings of perceived sadness given after sad video-clips were higher than those given after all the other video-clips s (all Zs ≥ 4.660; all *p*s ≤ 0.0001). Since session 3 results are similar to those of sessions 1 and 2, they are plotted in the Supplementary Information (**Fig.S2**).

### Subjective visceral experience

#### Capsule in the stomach (Session 1)

The Friedman ANOVA comparing the perceived visceral experience (i.e. gastric, cardiac, respiratory sensations and arousal) triggered by the five types of video-clips (varying for their content, i.e. disgusting; fearful; happy; sad; and neutral) when the capsule was in the stomach was statistically significant (χ^2^(19) = 323.399; *p* < 0.0001), suggesting that participants perceived different visceral sensations after observing the different content of the video-clips. Planned post-hoc Bonferroni-corrected Wilcoxon matched-pairs tests showed that *gastric sensations* were higher when participants were asked to observe disgusting and fearful videos. In particular, gastric sensations evoked by disgusting and fearful videos were significantly higher than those evoked by happy video-clips (all *Z*s ≥ 2.824; all *p*s ≤ 0.005). Gastric sensations evoked by disgusting video-clips were significantly higher than those evoked by neutral video-clips (*Z* = 2.777; *p* ≤ 0.005), while gastric sensations evoked by fearful video-clips were only marginally (considering the Bonferroni corrected significance threshold of 0.0125) higher than those evoked by neutral video-clips (*Z* = 2.375; *p* = 0.018). Gastric sensations evoked by sad video-clips were significantly higher only with respect to those evoked by happy video-clips (*Z* = 2.830; *p* ≤ 0.005). As to the *cardiac sensations*, we found that all the emotional video-clips evoked higher cardiac sensations with respect to the neutral video-clips (all Zs ≥ 3.295; all *p*s ≤ 0.001). As for the *respiratory sensations*, we found that they were maximally evoked by disgusting, fearful and sad videos (i.e. those inducing negative emotions). In particular, respiratory sensations evoked by disgusting, fearful and sad videos were significantly higher than those evoked by happy and neutral video-clips, (all *Z*s ≥ 3.036; all *p*s ≤ 0.002). Happy video-clips evoked higher respiratory sensations than neutral video-clips (*Z* = 2.515; *p* ≤ 0.012). Finally, we found that all the emotional video-clips evoked higher *feeling of arousal* with respect to the neutral video-clips, (all *Z*s ≥ 3.295; all *p*s ≤ 0.001). See **Fig.3** for a graphical representation of these results.

**Fig.3.**
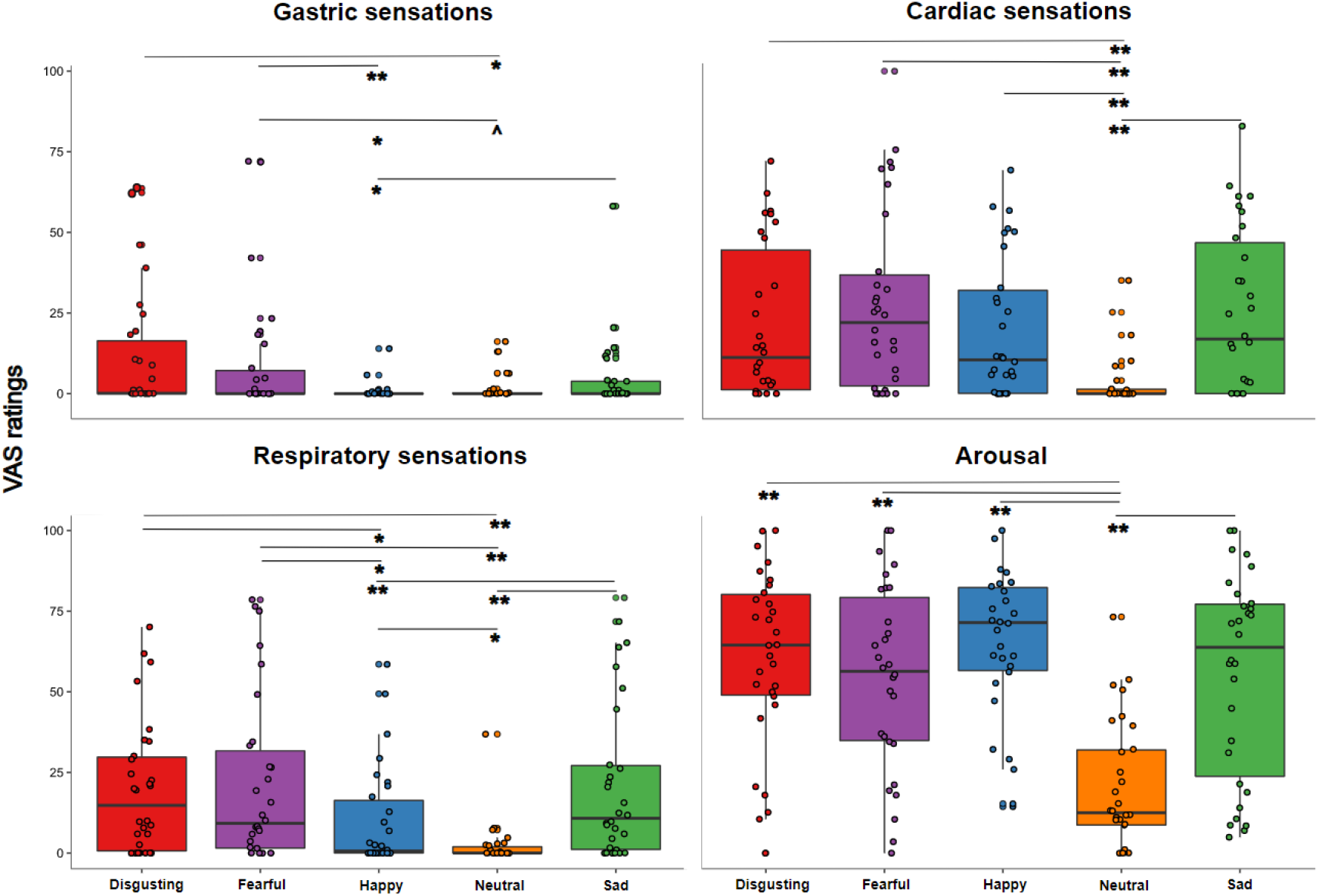
Visceral sensations results (pill in the stomach, session 1). Perceived visceral sensations (gastric, cardiac and respiratory) and arousal, measured using 0-100 visuo-analogue scale (VAS) ratings, as a function of the five categories of video-clips: disgusting, fearful, happy, sad, and neutral shown during the first session of this study (i.e., when the capsule was in the stomach). Significant differences refer to Bonferroni-corrected, Wilcoxon matched-pairs tests (threshold for multiple comparisons was set at 0.0125). ∧p = 0.018; * p < 0.01; ** p ≤ 0.001

#### Capsule in the small bowel (Session 2)

The Friedman ANOVA comparing the perceived visceral experience (i.e. gastric, cardiac, respiratory sensations and arousal) triggered by the five types of video-clips in the second session (namely when the capsule was in the small bowel) was statistically significant (χ^2^(19) = 289.385; *p* ≤ 0.0001), suggesting that participants perceived different sensations after observing the different content of the video-clips. Planned post-hoc Bonferroni-corrected Wilcoxon matched-pairs tests showed results akin to those found in the first session. Specifically, *gastric sensations* were higher when participants were asked to observe disgusting and fearful videos. In particular, gastric sensations evoked by disgusting videos were significantly higher than those evoked by all the other video-clips (all *Z*s ≥ 2.911; all *p*s ≤ 0.005), while gastric sensations evoked by fearful video-clips were higher than those evoked by happy and neutral video-clips (all *Z*s ≥ 2.58; all *p*s ≤ 0.01). As to the *cardiac sensations*, we found that all the emotional video-clips evoked higher cardiac sensations with respect to the neutral video-clips (all Zs ≥ 2.501; all *p*s ≤ 0.012) even though happy video-clips were only marginally significant considering the Bonferroni corrected significance threshold of 0.0125 (*Z* = 2.352.; *p* = 0.019). As for the *respiratory sensations*, we found that all the emotional video-clips but happy video-clips evoked higher respiratory sensations with respect to the neutral video-clips (all Zs ≥ 3.027; all *p*s ≤ 0.002). Moreover, respiratory sensations evoked by disgusting and sad videos were also significantly higher than those evoked by happy video-clips (all *Z*s ≥ 2.613; all *p*s ≤ 0.009). Finally, we found that all the emotional video-clips evoked higher *feeling of arousal* with respect to the neutral video-clips, (all *Z*s ≥ 4.28; all *p*s ≤ 0.0001). All the Wilcoxon matched-pairs comparisons that remained significant after Bonferroni-correction are shown in the Supplementary Information in **Fig.S3** to avoid redundancy.

#### Capsule in the large bowel (Session 3)

The Friedman ANOVA comparing the perceived visceral experience triggered by the five types of video-clips when the capsule was in the large bowel was statistically significant (χ^2^(19) = 286.999; *p* ≤ 0.0001), suggesting that participants perceived different sensations after observing the different content of the video-clips. *Gastric sensations* were higher when participants were asked to observe disgusting and fearful videos. In particular, gastric sensations evoked by disgusting videos were significantly higher than those evoked by all the other video-clips (all *Z*s ≥ 2.656; all *p*s ≤ 0.008), while gastric sensations evoked by fearful video-clips were higher than those evoked by neutral video-clips (*Z* ≥ 2.667; *p* ≤ 0.008). As to the *cardiac sensations*, we found that all the emotional video-clips evoked higher cardiac sensations with respect to the neutral video-clips (all Zs ≥ 3.419; all *p*s ≤ 0.001). As for the *respiratory sensations*, we found that all the emotional video-clips evoked higher respiratory sensations with respect to the neutral video-clips (all Zs ≥ 2.844; all *p*s ≤ 0.004). Moreover, respiratory sensations evoked by disgusting and sad videos were also significantly higher than those evoked by happy video-clips (all *Z*s ≥ 2.632; all *p*s ≤ 0.008). Finally, we found that all the emotional video-clips evoked higher *feeling of arousal* with respect to the neural video-clips, (all *Z*s ≥ 3.575; all *p*s ≤ 0.001). All the Wilcoxon matched-pairs comparisons that remained significant after Bonferroni-correction are shown in the Supplementary Information in **Fig.S4** to avoid redundancy.

### Gut markers of perceived emotions: pH, temperature, pressure and EGG peak frequency

#### Capsule in the stomach (Session 1)

Model 1 (see *Data analysis* session) had a marginal *R*^*2*^ = 0.51 and a conditional *R*^*2*^ = 0.60. Visual inspection of the plots did not reveal any obvious deviation from homoscedasticity. Residuals were not normally distributed according to Shapiro-Wilk normality test, but linear models are robust against violations of normality (Gelman and Hill, 2007). As for collinearity (tested by means of *vif* function of *car* package), all independent variables had a (GVIF^(1/(2*Df)))^2 < 10) except for pressure ((GVIF^(1/(2*Df)))^2 = 28.858) and pH ((GVIF^(1/(2*Df)))^2 = 17.986). Type III analysis of variance of Model 1 showed a statistically significant 2-way interaction between item (i.e. perceived disgust, fear, happiness and sadness) and gastric pH (*F* = 8.214, *p* < 0.0001, bootstrap p-value < 0.001, **Fig.S5**), suggesting that the emotional experience reported by participants on the VAS ratings, irrespective of the type of observed video clip, varied according to the pH of the stomach and the type of perceived emotion. Specifically, the follow-up post hoc simple slope analysis showed that the lower (i.e., more acidic) was the pH of the stomach, the more our participants reported to feel disgusted and afraid, while the higher the pH of their stomach (i.e., less acidic), the more they reported to feel happy. For each 1-unit decrease in stomach pH, a predicted 15.59 ± 5.3 points increase in VAS ratings of disgust was reported (*t* = −2.939, *p* = 0.003). Similarly, for each 1-unit decrease in stomach pH, there was a predicted 10.86 ± 5.3 points increase in VAS ratings of fear (*t* = −2.047, *p* = 0.041). On the other hand, for each 1-unit increase in stomach pH, there was a predicted 21.16 ± 5.3 points increase in VAS ratings of happiness (*t* = 3.988, *p* ≤ 0.001). The 2-way interaction was further defined by a 3-way interaction between video-clip content, item, and pH (*F* = 1.978, *p* = 0.025, bootstrap *p-value* = 0.026, **Fig.4**), suggesting that emotional experience reported by participants on the VAS ratings varied also as a function of the content of the projected video-clips, besides the pH of their stomach and the type of emotion perceived. Follow-up post hoc simple slopes analysis showed that the lower (i.e. more acidic) was the pH of the stomach, the more our participants reported to feel disgusted and afraid, after observing disgusting video-clips, while the higher was the pH of their stomach (i.e. less acidic) the more reported to feel happy. For each 1-unit decrease in stomach pH, there was a predicted 15.59 ± 5.3 points increase in VAS ratings of disgust (*t* = −2.939, *p* = 0.003, see the **first panel of Fig.4**). Similarly, for each 1-unit decrease in stomach pH, there was a predicted 10.86 ± 5.3 points increase in VAS ratings of fear (*t* = −2.047, *p* = 0.041, **see the first panel of Fig.4**). On the other hand, for each 1-unit increase in stomach pH, there was a predicted 21.16 ± 5.3 points increase in VAS ratings of happiness (*t* = 3.989, *p* ≤ 0.001, s**ee the first panel of Fig.4**). Simple slopes analysis also showed that the higher (i.e., more basic) was the pH of the stomach, the more our participants reported to feel happy after observing fearful video-clips. For each 1-unit increase in stomach pH, there was a predicted 7.51 ± 3.11 points increase in VAS ratings of happiness (*t* = 2.412, *p* ≤ 0.05, **see the second panel of Fig.4**). 3-way interactions with stomach pressure and temperature were not statistically significant (all *F*s ≥ 0.61; all *p*s ≤ 0.834). For a detailed description of the additional model results please refer to the Supplementary Information, please see **Table S1.A** in the Supplementary Information.

**Fig.4.**
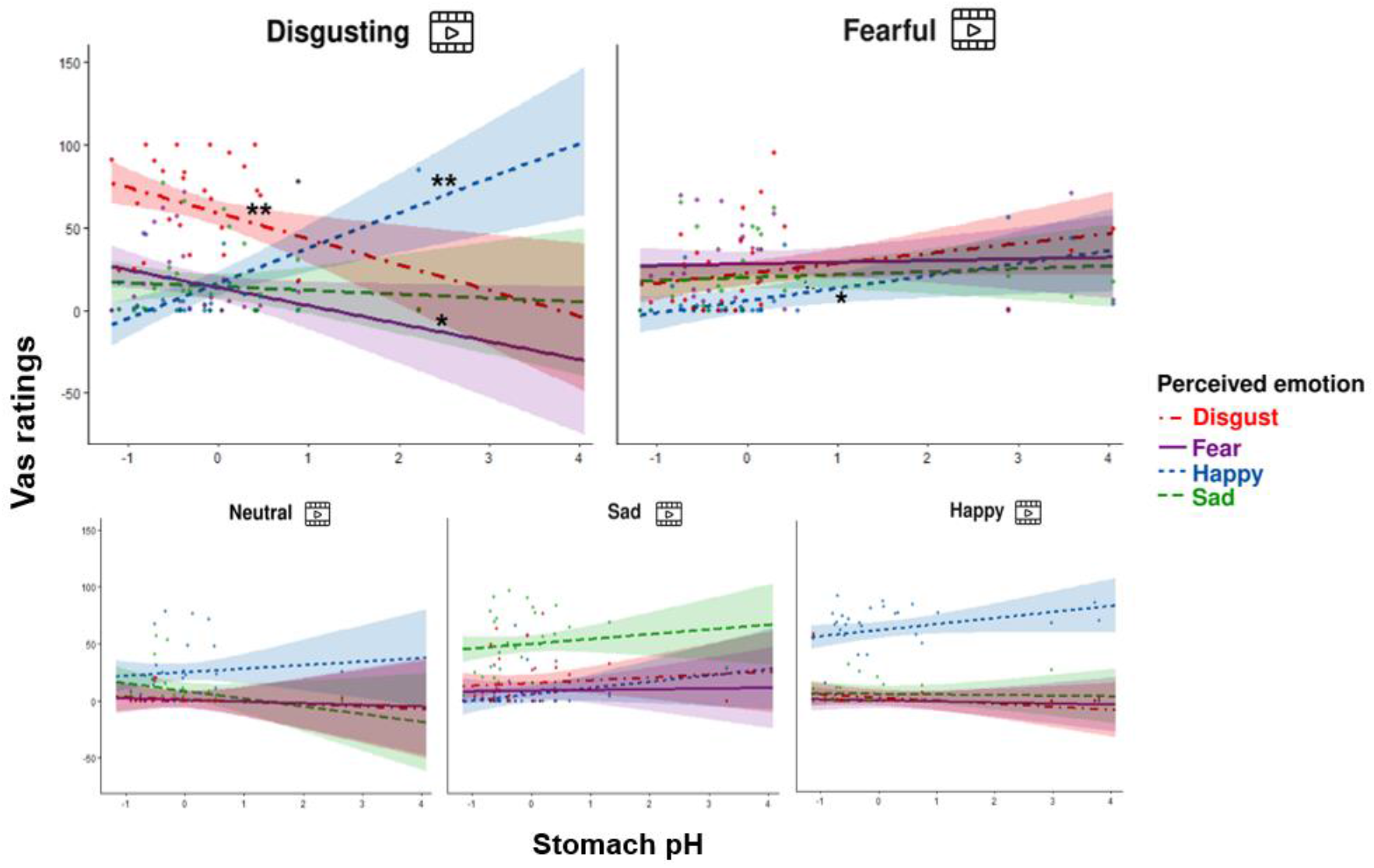
Stomach pH influences perceived emotions (3-way interaction between item, stomach pH and type of video-clip). Effects of stomach pH on perceived emotions (disgust, fear, happiness, and sadness) as a function of the five categories of video-clips (disgusting, fearful, happy, neutral, and sad). *p≤ 0.05; ** p≤ 0.01

With respect to the results of Model 4 (see above, *Data analysis* paragraph), i.e. the one containing the EGG peak frequency as independent variable, the model had a marginal R^2^ = 0.48 and a conditional R^2^ = 0.56. Visual inspection of the plots did not reveal any obvious deviation from homoscedasticity. Residuals were not normally distributed according to Shapiro-Wilk normality test, but linear models are robust against violations of normality (Gelman and Hill, 2007). As for collinearity (tested by means of *vif* function of *car* package), all independent variables had a (GVIF^(1/(2*Df)))^2 < 10). Type III analysis of variance of Model 4 showed a statistically significant 2-way interactions between video-clip content and item (F = 35.49, *p* < 0.001), however the 3-way interaction between video-clip content, item, and EGG peak frequency was not statistically significant (F = 0.57, *p* = 0.868), suggesting that emotional experience reported by participants on the VAS ratings was only influenced by the content of the projected video-clips, but not by the EGG peak frequency of their stomach. For a detailed and description of the model results please refer to **Table S2** of the Supplementary Information.

#### Capsule in the small bowel (Session 2)

Model 2 (see above, *Data analysis* procedure) did not report any significant 2- and 3-way interactions between video-clip content, items and small bowel pH, pressure, and temperature, suggesting that the emotional experience reported by participants on the VAS was only influenced by the content of the projected video-clips, but not by pH, pressure, or temperature recorded in the small bowel. For a detailed report of the model results please refer to **Table S3** of the Supplementary Information.

#### Capsule in the large bowel (Session 3)

Model 3 (see above, *Data analysis* procedure) did not report any significant 2- and 3-way interactions between video-clip content, items and small bowel pH, pressure, and temperature, suggesting that the emotional experience reported by participants on the VAS was only influenced by the content of the projected video-clips, but not by pH, pressure, or temperature recorded in the large bowel. For a detailed report of the model results please refer to **Table S4** of the Supplementary Information.

## Discussion

To explore the direct link between gastro-intestinal (GI) physiology and emotional experiences, we asked healthy male participants to observe a series of validated video-clips (Tettamanti et al., 2012) while electrical, chemical, and thermal signals originating from their GI system were recorded. Specifically, we induced disgust, fear, sadness, and happiness three times, when an inert ingestible pill that gauged pH, temperature and pressure was in the stomach, small and large bowel. When the pill was in the stomach we also recorded, via EGG, participants’ gastric myoelectric activity.

In addition to the objective GI measures, we collected participants’ subjective perception evoked by the different stimuli. Specifically, at the end of each emotional block, we asked participants to evaluate by means of 0-100 visuo-analogue scales (VAS) their perceived emotions, visceral sensations (gastric, cardiac, and respiratory) and arousal. In keeping with previous findings (Tettamanti et al., 2012), we found that the four emotional categories of the video-clips were clearly able to induce all the intended emotions (**Fig.2**). The effect was present in all the three experimental sessions, indicating that emotional induction was successful and no habituation over time occurred. It is also worth noting that the four types of emotional video-clips differed from the neutral ones which were perceived as less arousing (**Fig.3, bottom right panel**). These findings are consistent with previous literature suggesting that arousal is indeed an essential component of the emotional experience (Posner et al., 2005; Russell and Feldman Barrett, 1999) even if, unlike previous findings (Vianna et al., 2006; Vianna and Tranel, 2006), we did not find higher ratings of arousal triggered by negative vs. positive video-clips suggesting that the reported effects are genuinely associated with specific emotions.

Our results on the visceral sensations suggest that the subjective perception of different bodily signals, particularly of gastric feelings, varied across the different emotions. Specifically, gastric sensations were higher after observing fearful videos and maximal after observing disgusting videos, compared to the other emotional video-clips (**Fig.3, upper left panel**). Respiratory sensations, in turn, were higher after observation of disgusting, fearful and sad video-clips, compared to the happy and neutral ones (**Fig.3, bottom left panel**). Cardiac sensations instead did not differ between the emotional categories and were higher with respect to non-emotional stimuli (**Fig.3, upper right panel**). These results confirmed the idea that interoception (i.e. the awareness of physiological changes coming from internal organs) is a key component of emotional feelings. This is in line with theories of emotions (Barrett, 2017, 2014) that suggest a key role of bodily signals in the experience of emotions. However, the results also suggest that specific visceral signals uniquely characterize each emotional state, indicating that conceiving emotions as discrete (in line with somatic theories of emotions Damasio, 1999; Harrison et al., 2010; James, 1994) provide a clearer description of our data.

With respect to our main hypothesis – that internally and externally recorded GI markers are linked to emotions – we found that gastric physiology does play a significant role in emotional experiences. Specifically, the more acidic the pH recorded in participants’ stomach, the more they reported disgust and fear; the less acidic the gastric environment, the more participants reported happiness, independently from the observed category of video-clips (**Fig.S5**). Moreover, when participants were exposed to disgusting video-clips, more acidic gastric pH was associated with higher reports of disgust and fear, while less acidic gastric pH was associated with higher reports of happiness. Similarly, when participants were exposed to fearful video-clips, the less acidic was their pH, the more they reported to perceive happiness (**Fig.4**). Overall, these results were specific for the stomach and did not extend to the (small and large) bowel. Although it is true that the order was fixed, so that intestine data were always recorded after participants had already been exposed to a first round of video-clips when the capsule was in the stomach, at the subjective level (i.e. emotional and visceral VAS ratings) emotional stimuli did induce the same experience in all the three sessions. In the light of this evidence and of the literature, it seems that stomach physiology, and not a pure novelty effect triggered by the emotional stimuli, plays a specific and crucial role in subjective emotional experience.

Our findings about the link between pH acidity of the stomach and perceived emotions (disgust, fear, and happiness) are in line with the anecdotal reports described by Beaumont in 1833 (Beaumont, 1833) about Alexis St. Martin. He was a Canadian fur-trader with a gunshot-created permanent fistula, through which Beaumont could get access secretion samples from the patient’s stomach and report that when he was exposed to emotional experiences, mainly stress and anger, his gastric secretion changed color. Following these first inspiring but qualitative findings, very few studies explored changes in the human gastric milieu during higher-order cognitive and emotional processes. To the best of our knowledge, two single case studies conducted in the past century reported changes at the level of gastric acid output following anxiety induction procedure (Bennett and Venables, 1920) and a decrease in stomach motility and acid secretion during depression (Wolf and Wolff, 1947). Only after more than four decades, a somewhat systematic investigation showed that an acute psychological stress (i.e. caused by performing mental arithmetic and solving anagrams) not only increased arterial blood pressure and heart rate, but also stimulated gastric acid output recorded by means of a nasogastric tube) in healthy participants scoring high in impulsivity traits (Holtmann et al., 1990). Although in the present study we did not measure gastric secretion, which is a procedure that usually relies on invasive techniques (Ghosh et al., 2011), we show that stomach pH might be an important marker of emotional experience and that ingestible pills could be a non-invasive and sensitive tool to measure it.

As to the EGG data, we did not find any association between the peak frequency of the normogastric band (∼ 2-4 cpm) and the subjective experience of emotion. Future studies with longer emotional induction procedures are needed to better investigate if emotional video-clips might affect the other (bradygastric and tachigastric) bands. Unlike stomach pH, neither GI pressure nor temperature were associated with the emotional experience in the present study. It is important to underline that SmartPill capsules record temperature with a relatively low sample rate (every 20 s). Therefore, it is possible that different technologies with higher sample rates might be more sensitive to detect changes of endoluminal temperature triggered by the emotional experience.

Overall, our findings support somatic theories of emotions (Damasio, 1999; Harrison et al., 2010; James, 1994), suggesting that specific patterns of physiological signals are linked to unique emotional states (Stephens et al., 2010) and are in line with the behavioural and neural evidence highlighting the discrete structure of emotions (Lettieri et al., 2019; Stephens et al., 2010).

We acknowledge that our results, although novel, are merely correlational and restricted to a sample of male participants. To overcome these limitations, future studies involving females and investigating the causal nature of this link are needed. For example, manipulating the stomach pH during the emotional induction of basic, but also complex emotions, such as moral disgust (Rozin et al., 2009), would allow researchers to test the causality of this relationship. In particular, it would be interesting to see how emotions are experienced when classic anti-acids or proton pump inhibitors are administered or to see what happens to emotional experience after normalization of the gastric rhythm when an *anti-emetic* and a prokinetic agent, such as domperidone, is administered (Nord et al., 2021).

In conclusion, we believe that the present findings have the potential to open new avenues for studying the unexplored influences of the neurobiology of the gastro-intestinal system on typical and atypical emotional processes. Indeed, our approach may be adopted when studying the contribution of the gastric signals in people showing dysfunctional emotional processing like autism spectrum conditions and depression. Evidence shows that the above-mentioned conditions are characterized by comorbidities with GI problems (people with irritable bowel syndrome (IBS) are much more likely to develop depression than healthy controls (Shah et al., 2014), and autistic persons are more likely to develop IBS compared to controls (Kim et al., 2022)). Finally, our approach can be also adopted for studying emotional processes in conditions characterized by alterations of the gastro-intestinal physiology itself and its awareness, like persons with eating disorders (for a recent review see (Khalsa et al., 2022)).

## Materials and Methods

### Participants

To avoid the confounding effects played by sex both at level of emotional processing/experience (e.g. Kret and de Gelder, 2012) and of gastric physiology (e.g. Tolj et al., 2007) we choose to recruit only male participants to test a more homogeneous sample. Also, considering sex differences within emotional experience, testing men constitutes a conservative approach. We are well aware, though, that studies in women are necessary to generalize these findings and we are already working on this aspect.

Here, thirty-one healthy male participants (age: M = 24.42, S.D. = 2.8, range = 20–30 years) were recruited via the Social and Cognitive Neuroscience Laboratory volunteer database and through personal contacts to take part in this study. A preliminary structured questionnaire ensured they all were eligible to the experimental procedure and to the SmartPill™ (SmartPill Motility Testing System, Medtronic plc) capsule ingestion (see the SmartPill™ system paragraph for a detailed description of this device). Exclusion criteria included diagnosis of psychiatric, neurological, or swallowing disorders; gastric bezoars; history of any abdominal/pelvic surgery within the previous three months; suspected or known strictures, fistulas, or physiological/mechanical obstruction within the gastro-intestinal tract; dysphagia to food or pills; Crohn’s disease or diverticulitis; body mass index ≥ 40; age < 18 years; cardiac pacemakers or defibrillators; assumption of any medication or substance that could interfere with pH values and gastro-intestinal motility (Saad, 2016), such as tobacco (within 8 h prior to capsule ingestion), antacids and alcohol (within 24 h), laxatives (within 48 h); antihistamines, prokinetics, antiemetics, anticholinergics, antidiarrheals, narcotic analgesics, non-steroidal anti-inflammatories (within 72 h), or proton pump inhibitors (within seven days). The protocol of this study was approved by the local Institutional Review Board (Fondazione Santa Lucia ethics committee, Protocol CE/PROG.636) and was performed in accordance with the Declaration of Helsinki (1991). All participants were naïve to the purpose of the experiment, provided their written informed consent before starting the experimental procedure, and received a reimbursement for their participation at the end of the study. The same day of the experiment, they also took part in two additional studies, each with different aims and different hypotheses.

### SmartPill™ system

The SmartPill™ system (SmartPill Motility Testing System, Medtronic plc) was originally approved by the US Food and Drug Administration for the evaluation of suspected gastroparesis and colonic transit in patients with chronic idiopathic constipation (Aburub et al., 2018). In a position paper, the American and European Neurogastroenterology and Motility Societies indicated that this system is useful in the assessment of clinical disorders affecting the gastro-intestinal tract (Rao et al., 2011).

The SmartPill™ system consists of a wireless, single-use, ingestible capsule; a receiver; a docking station; and a software (i.e., MotiliGI) installed on a dedicated laptop (**Fig.2, panel A**). The ingestible pill is a cylindrical polyurethane capsule, 26.8 mm long, 11.7 mm wide, and it weighs 4.5g (Hasler, 2014). It houses a pH sensor with a range of 1–9 units (accuracy of ± 0.5 units); a solid state temperature sensor with a range of 20–42°C (accuracy of ± 1°C), a solid-state pressure sensor with a range of 0–350 mmHg (accuracy of ±5 mmHg for values < 100 mmHg and accuracy of ± 10% for values > 100 mmHg), two silver oxide batteries (duration > 5 days), and a transmitter (broadcast frequency: 434.2 MHz). Prior to ingestion, the capsule is activated through a magnetic fixture and the pH sensor is calibrated through a buffer solution. After ingestion, the capsule starts recording intraluminal pressure, pH, and temperature and sending these data to the external radio receiver that participants wear around their waist. For the first 24 h, temperature data are transmitted every 20 s, pressure every 0.5 s, and pH every 5 s; after the first day, sampling frequencies are halved. By means of the MotiliGI software (Medtronic), pH, temperature and pressure data can be graphically displayed in real time or reviewed after the pill expulsion. Combining pH and temperature data, MotiliGI identifies the specific region of the gastro-intestinal tract in which the pill is located. The arrival of the capsule in the stomach is indicated by a rapid rise in temperature from ambient to body temperature and by a drop in pH (reflecting passage into the acidic environment of the stomach). The arrival of the capsule in the small intestine through the pylorus is instead indicated by an abrupt increase of ≥ 2 pH units from the gastric baseline. Likewise, a subsequent gradual decrease of ≥ 1 pH unit for at least 10 consecutive minutes is considered as a sign that the pill left the small intestine and entered the large intestine. If the pH decrease cannot be observed, the MotiliGI software relies on pressure data to mark the transition between small and large bowel. The expulsion of the capsule is defined by a drastic drop in temperature followed by loss in recorded signal after the subject defecated. In our sample, 30 out of 31 subjects displayed a prototypical pattern of pH increase and decrease, thus indicating when the pill was in each of the three areas of interest. For the remaining subject, the software was still able to localize the gastro-intestinal regions that the pill went through based on temperature, pressure, and time data.

### Electrogastrography (EGG)

During the first session of this study (see the *Experimental procedure* paragraph for a detailed description), besides gastro-intestinal pH, temperature, and pressure, we also recorded participants’ surface gastric myoelectric activity by placing three cutaneous electrodes on their abdominal skin over the stomach (**Fig.5, panel B**). Electrogastrography (EGG) is a non-invasive technique that allows the recording of i) the slow (∼0.05 Hz) electrical gastric rhythm, constantly generated in the stomach wall within the so-called “dominant pacemaker” area located in the greater curvature of the mid/upper corpus; ii) the more transient smooth muscle activity inducing gastric peristaltic contractions (see (Wolpert et al. (, 2020) for an extensive review). Here, we used a 1-channel EGG bipolar montage that consists of three pre-gelled disposable Ag/AgCl electrodes placed in a standardized position (Yin and Chen, 2013) and attached to a PowerLab data acquisition device (ADInstrument Ltd). Specifically, the first recording electrode was placed halfway between participants’ xiphoid and their umbilicus, the second 5 cm up and 5 cm to the left of the first (taking the left side of participants as a reference), and the ground electrode was placed on the left costal margin (see the **left part of Fig.5**).

**Fig.5.**
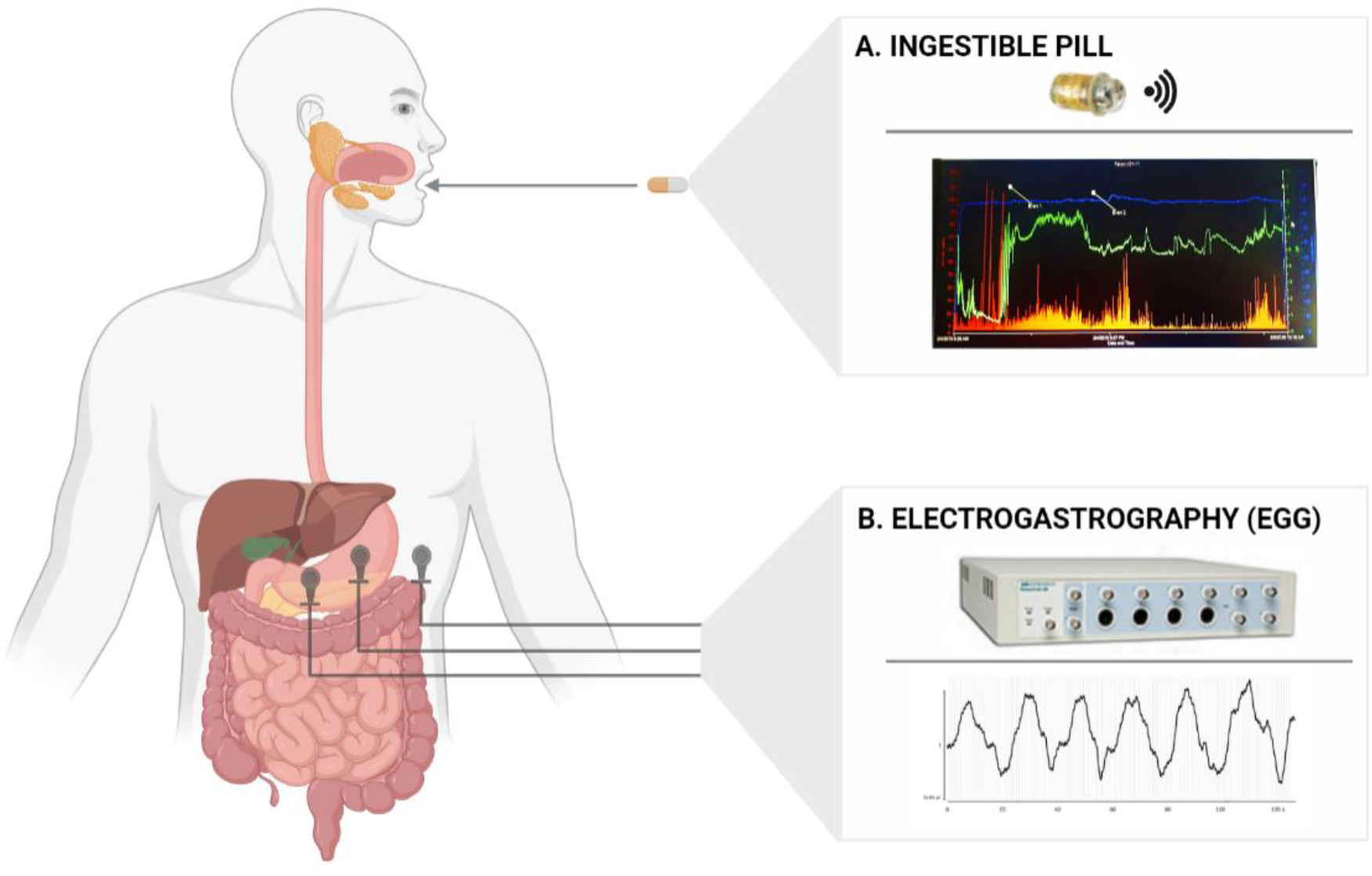
Gastro-intestinal (GI) markers collected in the present study. Panel A shows the SmartPill capsule used to record temperature, pH, and pressure of the gut and the intestine as well as a typical graph created by the dedicated MotiliGI software, plotting temperature (blue line), pH (green line), and pressure (orange bars) across the entire the gastro-intestinal tract. Panel B shows the device (PowerLab, ADInstruments Ltd) used to record electrogastrography (EGG) and a typical gastric myoelectric activity of a participant. The 1-channel bipolar electrode montage for the EGG recording is displayed on the left side of the figure. This figure was designed with BioRender.com and edited with Adobe Photoshop® 7.

### Emotional induction task

An illustration of the emotional induction procedure adopted in this study is provided in **Fig.1**. Participants were asked to observe four blocks of a series of emotional video-clips with different emotional content. Each block aimed at triggering a specific basic emotion, namely disgust, fear, happiness, and sadness. In addition to the four blocks, participants also observed one block containing a series of neutral video-clips (control condition). The five blocks were presented in randomized order across participants and contained 24 emotional video-clips each. Video-clips were edited color soundless film excerpts lasting ∼9 s each. The order of video-clips within each block was randomized. All the 115 video-clips were selected and validated by Tettamanti and collaborators (Tettamanti et al., 2012). Clips showing mutilations or contamination scenarios (e.g. getting in touch with offensive or infective agents) triggered disgust; clips portraying human or animal aggressions and dangerous situations (e.g. falling or drowning) elicited fear; clips depicting amusement scenes, significant achievements or meaningful human interactions (e.g. a meeting between lovers) triggered happiness; clips containing scenarios of death, loss or sickness, (e.g. someone holding their beloved dead) triggered sadness; finally, clips with scenarios of routine human actions and activities (e.g. working or speaking) did not contain any specific emotion and therefore were considered neutral. For a detailed description of the stimuli (and their validation) please refer to the original study (Tettamanti et al., 2012) and to the Supplementary Information (**Table S5**), in which a brief description of the content of each of the video-clips employed in our study is provided. At the end of each block, participants answered an 8-item questionnaire relative to their emotional and visceral feelings induced by the observation of the video-clips. Four items measured the *emotional experience* in terms of perceived intensity of each emotion, three additional items measured the *visceral experience* in terms of perceived intensity of each visceral feeling and one item measured the perceived *arousal*. Items were presented in a randomized order and participants provided their responses by clicking with the mouse on separate 0 (“Not at all”) - 100 (“Very much”) visuo-analogue scales (VAS). To avoid carryover effects induced by each category of the emotional video-clips, blocks were interspersed with two-minutes washout pauses in which participants simply rested. Instructions, stimuli presentation and collection of responses concerning the emotional induction task were handled by a custom MATLAB algorithm. **Table 1** shows the complete list of the questionnaire items following each block of video-clips.

**Table 1.** Eight-item questionnaire. English translation and original Italian text of the 8-items questionnaire probing the participants’ emotions, visceral feelings, and arousal after each block of video clips. Ratings were provided along 0-100 VAS scales.

| Construct | Category | Item (English) | Item (Original language: Italian) |
| --- | --- | --- | --- |
| ***Emotional experience*** | Perceived disgust | How much disgust did you feel? | Quanto disgusto hai provato? |
| ***Emotional experience*** | Perceived fear | How much fear did you feel? | Quanta paura hai provato? |
| ***Emotional experience*** | Perceived happiness | How much happiness did you feel? | Quanta felicità hai provato? |
| **Emotional experience** | Perceived sadness | How much sadness did you feel? | Quanta tristezza hai provato? |
| **Visceral experience** | Perceived gastric sensations | Did you feel retching, stomach contractions, stomach cramps or other gastro-intestinal sensations? | Hai provato conati di vomito, contrazioni gastriche, crampi allo stomaco o altre sensazioni gastrointestinali? |
| **Visceral experience** | Perceived cardiac sensations | Did you feel your heart change (i.e. accelerate/slow down or beat)? | Hai sentito il tuo cuore cambiare (accelerare/rallentare/palpitare)? |
| **Visceral experience** | Perceived breathing sensations | Did you feel your breath change (heavy breath /holding breath/lack of air?) | Hai sentito il tuo respiro cambiare (respirare affannosamente/trattenere il respiro/mancanza d'aria)? |
| **Arousal** | Perceived arousal | Did watching the clips bored/aroused you? | Vedere i video ti ha annoiato/coinvolto? |

### Experimental procedure

Testing sessions began with an overnight fast, the avoidance of tobacco and alcohol, and the discontinuation of medications potentially altering gastric pH and gastro-intestinal motility (see above). Participants read, filled, and signed the informed consent form. After verifying that all the requirements for the participation were met, participants ate a standardized ∼260 kcal breakfast consisting of egg whites (120 g), two slices of bread and jam (30 g) to make sure that gastro-intestinal transit times of the SmartPill capsule were not affected by meal variability. Meanwhile, in a dedicated room, we activated the capsule through a magnetic fixture and calibrated the capsule pH sensor (see Materials above). After pH calibration was complete, the pill started transmitting data to the radio receiver. All but one pH calibrations were performed correctly, therefore pH data from one participant were not considered in the statistical analyses since they were not reliable (i.e. pH calibration was not correctly performed). Data recorded through the capsule came with a relative timestamp indicating the number of seconds elapsed from calibration. Since SmartPill data were not associated with an absolute time frame, we synchronized each calibration with an external clock that provided us with the required absolute time frame (see *Data analysis* session for a detailed description of the data pre-processing). At that point, participants swallowed the SmartPill capsule while drinking a glass of water (120 ml). A medical doctor supervised the ingestion procedure to help in case of swallowing problems. All participants ingested the pill without any trouble. After the ingestion, they fastened the receiver around their belt and lay supine on a deck chair. The experimenter cleaned the skin corresponding to the position of the EGG electrodes and afterwards attached them according to a 1-channel bipolar montage (Yin and Chen, 2013) (see **Fig.5**). After checking that a highly acidic pH (∼1-2) was recorded by the capsule, indicating that it reached the stomach, a 15-minute resting-state SmartPill capsule/EGG baseline session started. During this 15-minute resting-state session, participants were left alone in the room and instructed to relax while keeping their eyes open. At the end of the baseline session, participants underwent the emotional task during which they observed five blocks of film excerpts while stomach pH, temperature, pressure, and EGG signal were recorded. After each block, participants answered the 8-item questionnaire relative to their subjective emotional and visceral experiences triggered by the video-clips. Participants repeated the emotional task two more times, namely when the SmartPill capsule reached the small intestine (usually after 2 – 5 hours from the capsule ingestion) and when it reached the large intestine (usually after 2 – 6 hours from the stomach-small bowel transition). After around 6 hours from the beginning of the experimental session, participants were provided with a meal and *ad libitum* water. After 8 hours, they were also allowed to smoke. Between the experimental sessions they were free to work, study, or rest, although they were asked to avoid strenuous physical exercise and alcohol consumption. At the end of the last experimental session, i.e. at the end of the emotional task and while the capsule was in the large bowel, participants could leave the laboratory. They were asked to keep the receiver with them until they noticed that after defecation data transmission stopped due to pill expulsion, an event that ordinarily happens after 10-59 hours from ingestion (Saad, 2016). After the expulsion, participants came back in the lab to return the receiver.

## Data analysis

Details on SmartPill and EEG data pre-processing can be found in the Supplementary Information.

## Statistical data analyses

To exclude habituation effects, we tested whether the five blocks of video-clips triggered different subjective emotional and visceral experiences in our participants in all the three experimental sessions. To do so, separate Friedman ANOVAs followed by Bonferroni-corrected Wilcoxon matched-pairs tests (*p* ≤ 0.0125 was considered as significance threshold) were performed on the VAS ratings relative to the emotional and visceral experience triggered by the different types of video-clips. A non-parametrical approach was used because in several conditions VAS ratings were not normally distributed according to the Kolmogorov–Smirnov (K-S) tests and Skewness and Kurtosis z-scores, as suggested by Field (2009). Then, we estimated the impact of gastro-intestinal pH, temperature, and pressure (as measured by the SmartPill capsule) on the emotional experience (as reflected in the VAS ratings). To do so, three separate mixed-effects models were run, one for each GI region (stomach, small bowel, and large bowel). When statistically significant main or interaction effects were found, robust p-values were bootstrapped through the *mixed* function of the *afex* package (Singmann et al., 2015). Post-hoc simple slope analyses of the statistically significant interactions were run using the *sim_slopes* function of the *interaction* package (Long, 2022). A fourth mixed-effect model was run to estimate the impact of EGG peak frequency over the emotional experience triggered by the video-clips. The first three models had as dependent variable the VAS ratings relative to the emotional experience questionnaire, while fixed factors included *emotion*, i.e. the emotional content triggered by the video-clips (five levels: disgust, fear, happiness, sadness, neutral); *item*, i.e. the emotion perceived after observing the video-clip (four levels: perceived disgust, fear, happiness and sadness); and the block-specific *pH* (continuous), *pressure* (continuous), and *temperature* (continuous) recorded while participants observed the video-clips. pH, temperature, and pressure data were centred and scaled before undergoing further analysis. The fourth (EGG) model was similar to the other three, since it featured *emotion* and *item* as fixed factors, but had the block-specific (centred and scaled) *EGG peak frequencies* as continuous fixed factor. All the four models included by-subject intercepts as random effects.

Significant main or interaction effects were replicated if the random slope of the *emotion* was included as random effect in the mixed models. Moreover, to reduce the probability to describe false positives due to multiple comparisons, we used the *emtrends* function of the *emmeans* package (Lenth et al., 2019) to perform post hoc tests on the more conservative model (the one with the condition as random slope) with Bonferroni-adjusted *p* values (see Table S1.B contained in the supplementary materials for the detailed results). Due to the fact that models with random slopes yielded a boundary (singular) fit we decided to report the results coming from the less complex, intercept-only models in the main text. For the detailed description of the models and the software/packages used, please see the Supplementary Information.

## Data Availability

Data collected in this study are available at the OSF space dedicated to this project (https://osf.io/uq4t3/?view_only=ac0f717460e447c1be9df024dc3a2304). Any additional information or material requests can be addressed to Giuseppina Porciello or Salvatore Maria Aglioti.

## Author Contributions

Conceptualization and methodology: GP, AM, MSP, SMA. Experiment scripts and data pre-processing: AM, GP. Statistical analyses: GP, AM, MSP. Funding acquisition: SMA, MSP, GP. Resources and supervision: SMA. Visualization and writing – original draft: GP. Writing – review and editing: GP, AM, MSP, SMA.

## Acknowledgments

We thank Prof. Marco Tettamanti for sharing the full set of emotional stimuli, Danila Cosenza for medical assistance, Maurizio Molisso for standardized meal preparation, and our brave volunteers for their participation. The support of the Institut d’études avancées de Paris to SMA is gratefully acknowledged. This research was supported by ERC Advanced Grant 789058 (eHonesty) to SMA; by Regione Lazio grant A0375-2020-36612 to SMA and MSP; and by SEED PNR21 to GP.

