## Supplementary materials for "Deep-body feelings: ingestible pills reveal gastric correlates of emotions"

**Classification:** Neuroscience

**Keywords:** Emotions; Visceral sensations; Ingestible pills; pH; Gut; Interoception

### Supporting Information Text

#### Supplementary Information about data analysis

#### SmartPill data pre-processing

Once the pill was expelled and the receiver arrived in the lab, SmartPill data of each participant were uploaded and visualized through the MotiliGI software. Stomach, small bowel, large bowel, and whole gut transit times of the capsule (Lee et al., 2014) were calculated in order to estimate possible anomalies. All but one participant showed regular transit times. The remaining subject had an abnormal, i.e. too long large bowel transit time (> 59 h (Saad, 2016)). Consequently, his data were discarded from the statistical analyses. A custom MATLAB algorithm converted relative timestamps recorded by the SmartPill capsule in absolute times, so that each event (e.g. beginning and end of each experimental block of the emotional task) was paired to a definite hh:mm:ss:ms string.

#### EGG data pre-processing

Raw EGG signals were visually inspected on LabChart (data analysis software, ADInstruments Ltd) to remove artifacts due to body movements. A 0.016-0.15 Hz bandpass filter removed pink noise and unwanted higher frequencies associated with cardiac, respiratory, and small bowel activity. For each emotional block, we calculated the individual EGG peak frequency, namely the maximum periodogram peak found in the 'normogastric' range, i.e. the range of frequencies that is compatible with the number of stomach contractions in healthy individuals (0.033–0.066 Hz ~ 2-4 cycles per minute, cpm (Wolpert et al., 2020); EGG spectral density was computed using Welch's method on 200 s time windows with 150 s overlap (Rebollo et al., 2018). EGG analysis was performed with BrainVision Analyzer (Brain Products GmbH) and the MATLAB FieldTrip toolbox (Oostenveld et al., 2011).

#### Mixed model analysis

Mixed models were specified as follows.

**Model 1 (SmartPill data, stomach)**

*VAS* ratings *~ video-clip content * item * (ph + pressure + temperature) + (1 | subject), data = stomach_data, control = lmerControl(optimizer = "nloptwrap", calc.deriv= FALSE))*

**Model 2 (SmartPill data, small bowel)**

*VAS* ratings *~ video-clip content * item * (ph + pressure + temperature) + (1 | subject), data = smallbowel_data, control = lmerControl(optimizer = "nloptwrap", calc.deriv= FALSE))*

**Model 3 (SmartPill data, large bowel)**

*VAS* ratings ~ *video-clip content * item * (ph + pressure + temperature) + (1 | subject), data = largebowel_data, control = lmerControl(optimizer = "nloptwrap", calc.deriv= FALSE))*

**Model 4 (EGG data, stomach)**

*VAS* ratings ~ *video-clip content * item * egg peak frequency + (1 | subject), data = egg_data, control = lmerControl(optimizer = "nloptwrap", calc.deriv= FALSE))*

**Statistical softwares and packages**

Friedman ANOVAs followed by Bonferroni-corrected Wilcoxon matched-pairs tests were run using Statistica 7 software.

The remaining analyses were performed with R Studio. Specifically, we used the *lme4* package (Bates, D., Maechler, M., Bolker, B., & Walker, 2014) to perform linear mixed-effects analyses; the *lmerTest* package (Kuznetsova, 2017) to extract p-values through a Type III analysis of variance with Satterthwaite's method; the *mixed* function of the *afex* package (Singmann et al., 2015) to compute robust p-values with bootstrap method; the *interactions* (Long, 2019) and the *emmeans* (Length et al., 2019) packages to perform post-hoc simple slopes analysis and plots when significant interactions were found; and the *ggplots2* package (Wickham, 2016) to perform boxplots of the emotional, visceral and arousal experience. The standard assumptions and requirements of mixed models (linearity, homoscedasticity, absence of collinearity, and normality of residuals) were assessed through visual inspection of residual plots, the *shapiro.test* function and the *vif* function of the *car* package (Fox and Weisberg, 2019). The absence of singularity of the model was instead checked via the *check_singularity* function of the *performance* package (Lüdecke et al., 2021). Finally, the percentage of variance explained by each mixed-effects model was computed through the *r.squaredGLMM* function of Kamil Bartoń's *MuMIn* package.

**Supplementary results**

**Gut markers of perceived emotions: pH, temperature, pressure and EGG peak frequency**

**Capsule in the stomach (Session 1)**

Type III analysis of variance of Model 1 showed a statistically significant 2-way interaction between item and gastric pH, suggesting that emotional experience reported by participants on the VAS ratings varied according to the pH of the stomach and the type of perceived emotion, irrespectively of the type of observed video clip. The follow-up post hoc simple slope analysis showed that the lower (i.e., more acidic) was the pH of the stomach, the more our participants reported feeling of disgust and fear, while the higher was the pH of their stomach (i.e., less acidic) the more they reported feelings of happiness, see **Fig. S5** for a graphical representation of these effects. For the detailed description of Model 1 results refer to **Table S1** below.

**Capsule in the small bowel (Session 2)**

Model 2 (see the main text for data analysis procedure) did not show any problem of singular fit, had a marginal $R^2$ = 0.57 and a conditional $R^2$ = 0.66. Visual inspection of the plots did not reveal any obvious deviation from homoscedasticity. Residuals were not normally distributed according to Shapiro-Wilk normality test, but linear models are robust against violations of normality (Gelman and Hill, 2007). As for collinearity, all independent variables had a GVIF^(1/(2*Df)))^2 < 10.

Type III analysis of variance of Model 2 showed a statistically significant 2-way interaction between video-clip content and item ($F$ = 65.29, $p$ < 0.005), but no statistically significant 3-way interactions between video-clip content, item, and small bowel pH ($F$ = 0.60, $p$ = 0.84), or video-clip content, item, and small bowel pressure ($F$ = 1.44, $p$ = 0.14), or video-clip content, item, and small bowel temperature ($F$ = 1.65, $p$ = 0.07). This pattern of results suggests that emotional experience reported by participants on the VAS was only influenced by the content of the projected video-clips and the item type, but not by pH, pressure, or temperature recorded in the small bowel. For a detailed report of the model results please refer to the **Table S4.**

**Capsule in the large bowel (Session 3)**

Model 3 did not show any problem of singular fit, had a marginal $R^2$ = 0.45 and a conditional $R^2$ = 0.58. Visual inspection of the plots did not reveal any obvious deviation from homoscedasticity. Residuals were not normally distributed according to Shapiro-Wilk normality test, but linear models are robust against violations of normality (Gelman and Hill, 2007). As for collinearity, all independent variables had a GVIF^(1/(2*Df)))^2 < 10.

Type III analysis of variance of Model 3 showed a statistically significant 2-way interaction between video-clip content and item ($F$ = 42.46, $p$ < 0.005), but any statistically significant 3-way interactions between video-clip content, item and large bowel pH ($F$ = 0.61, $p$ = 0.83), or video-clip content, item and large bowel pressure ($F$ = 0.4, $p$ = 0.96), or video-clip content, item and large bowel temperature ($F$ = 0.56, $p$ = 0.88), suggesting that emotional experience reported by participants on the VAS was only influenced by the content of the projected video-clips and the item type, but not by pH, pressure, or temperature recorded in the large bowel. For a detailed report of the model results please refer to **Table S5**.

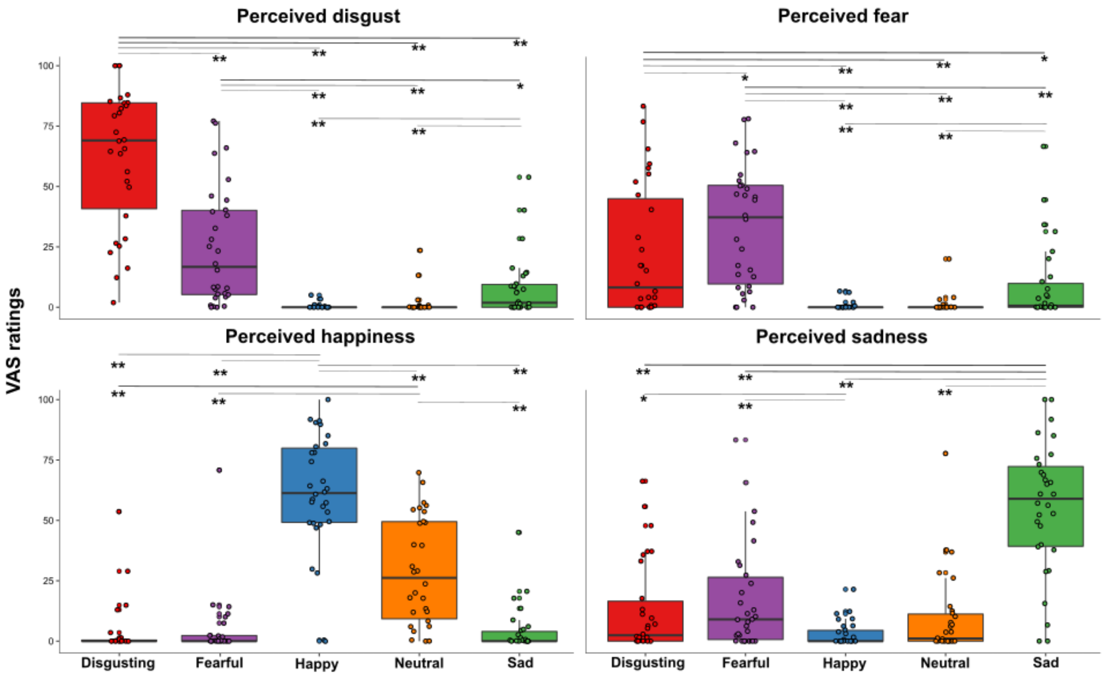


**Fig.S1 Perceived emotions results (pill in the small bowel, session 2).** Perceived emotions (disgust, fear, happiness, and sadness) measured using 0-100 visuo-analogue scale (VAS) ratings, as a function of the five categories of video-clips (disgusting, fearful, happy, sad, and neutral) shown during the second session of this study (i.e., when the capsule was in the small bowel). * $p \leq 0.05$; ** $p \leq 0.01$.

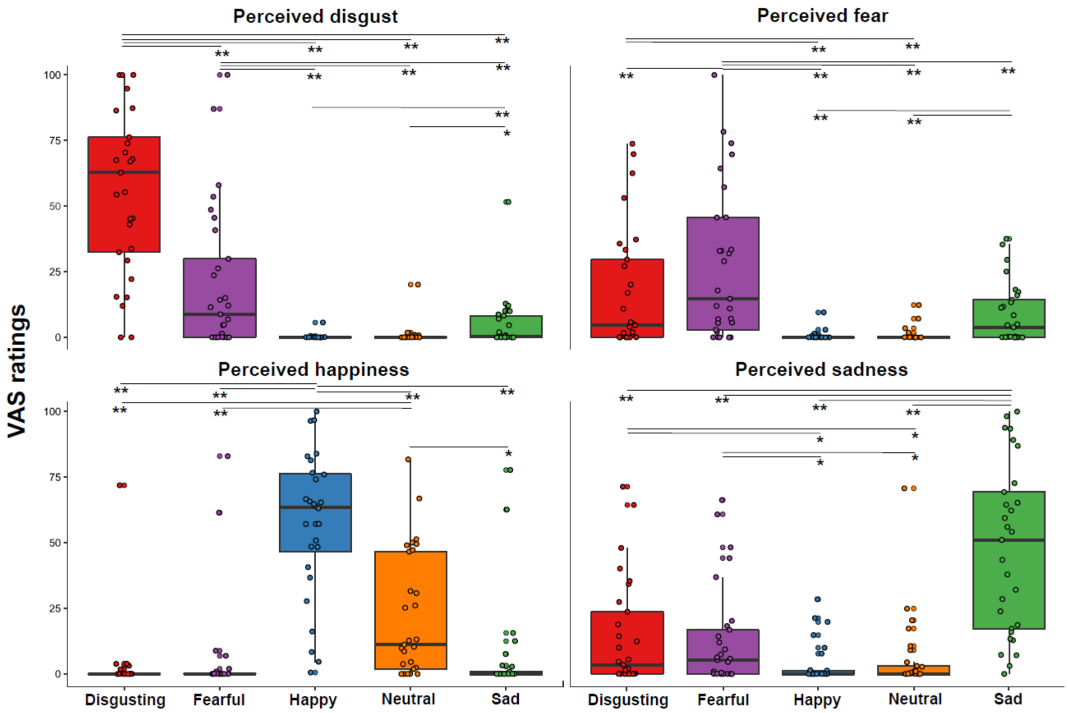


**Fig.S2 Perceived emotions results (pill in the large bowel, session 3).** Perceived emotions (disgust, fear, happiness, and sadness) measured using 0-100 visuo-analogue scale (VAS) ratings, as a function of the five categories of video-clips (disgusting, fearful, happy, sad, and neutral) shown during the third session of this study (i.e., when the capsule was in the large bowel). * $p \leq 0.05$; ** $p \leq 0.01$.

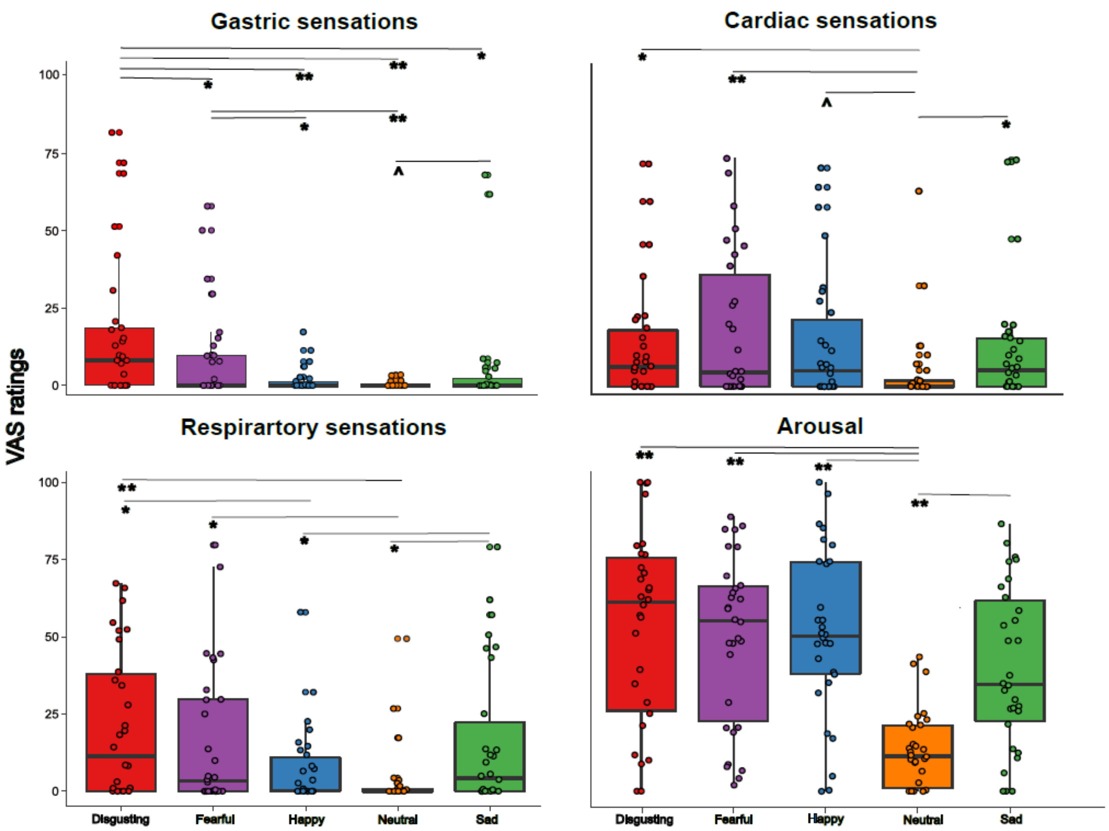


**Fig.S3 Visceral sensations results (pill in the small bowel, session 2).** Perceived visceral sensations (gastric, cardiac and respiratory) and arousal, measured using 0-100 visuo-analogue scale (VAS) ratings, as a function of the five categories of video-clips (disgusting, fearful, happy, sad, and neutral) shown during the second session of this study (i.e., when the capsule was in the small bowel). ^ $p = 0.019$; * $p \leq 0.01$; ** $p \leq 0.001$.

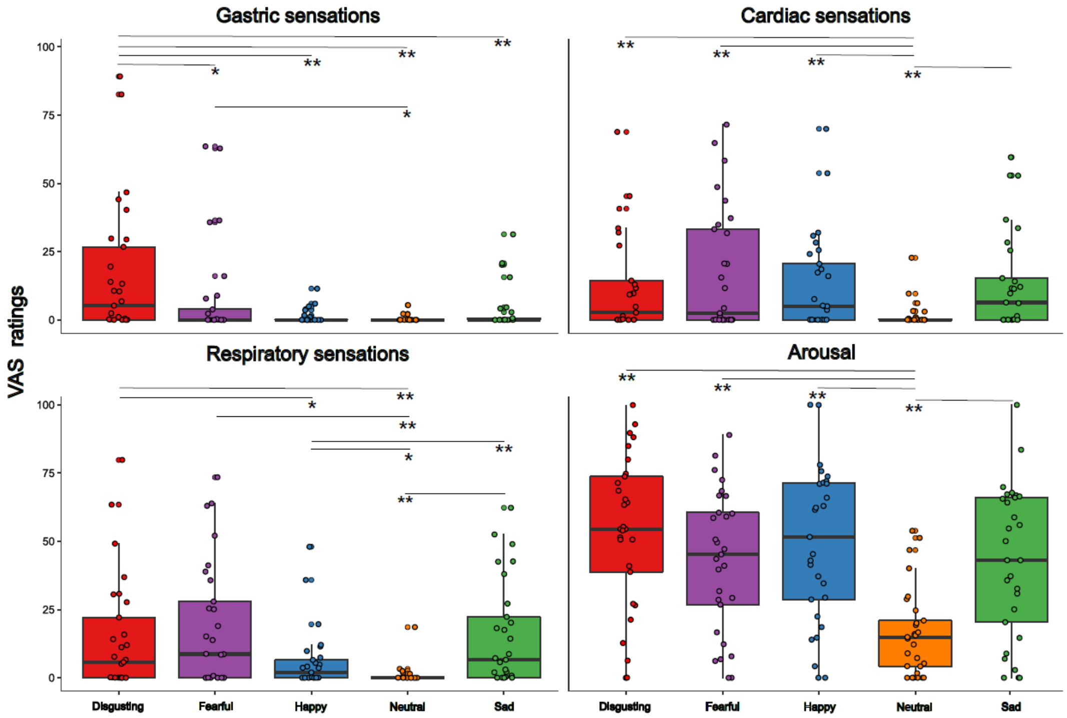


**Fig.S4 Visceral sensations results (pill in the large bowel, session 3).** Perceived visceral sensations (gastric, cardiac and respiratory) and arousal, measured using 0-100 visuo-analogue scale (VAS) ratings, as a function of the five categories of video-clips: disgusting, fearful, happy, sad, and neutral shown during the third session of this study (i.e., when the capsule was in the large bowel). ^ $p = 0.018$; * $p \leq 0.05$; ** $p \leq 0.01$.

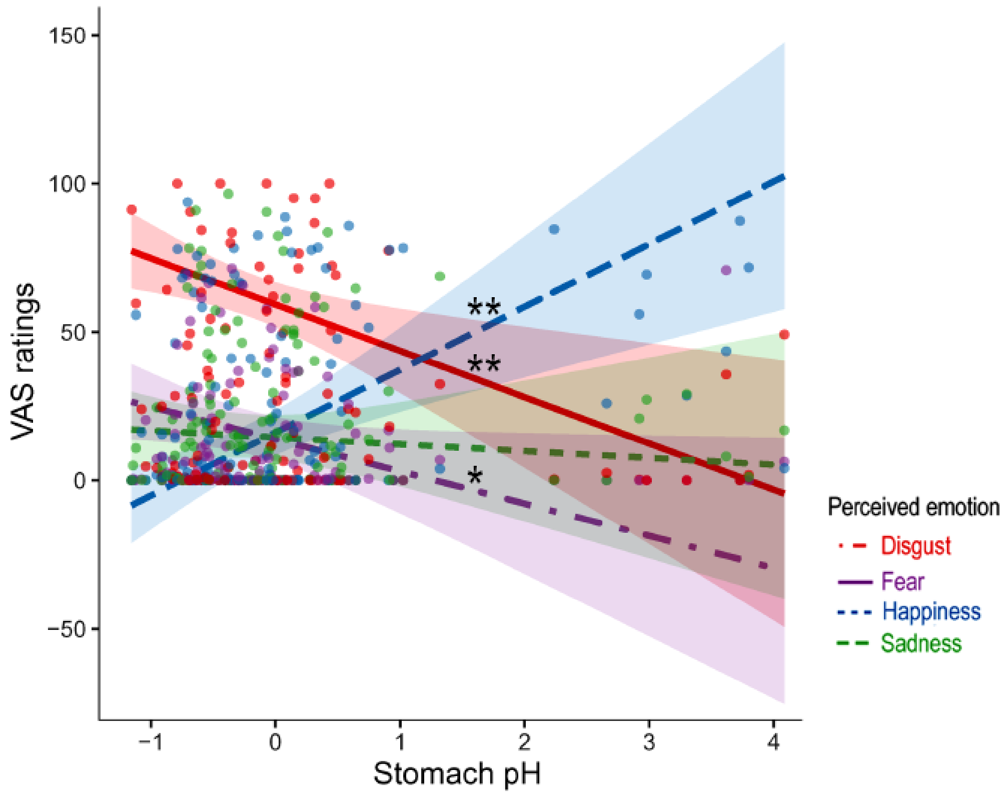


**Fig.S5 Stomach pH influences perceived emotions (2-way interaction between item and gastric pH).** Effects of stomach pH on perceived emotions (disgust, fear, happiness and sadness). * $p \leq 0.05$; ** $p \leq 0.01$

**Table S1**

**S1. A Results of Model 1 (pill in the stomach, session 1).** Type III analysis of variance table with Satterthwaite's method.

| | Sum Sq | Mean Sq | NumDF | DenDF | F value | P |
|---|---|---|---|---|---|---|
| **Video-clip content** | 14864 | 3715.9 | 4 | 460.33 | 11.8690 | 3.475e-09 *** |
| **Item** | 12406 | 4135.2 | 3 | 453.84 | 13.2081 | 2.785e-08 *** |
| **(Stomach) Pressure** | 1321 | 1320.6 | 1 | 286.71 | 4.2182 | 0.04090 * |
| **(Stomach) pH** | 80 | 79.7 | 1 | 96.07 | 0.2545 | 0.61511 |
| **(Stomach) T** | 314 | 314.3 | 1 | 31.69 | 1.0039 | 0.32397 |
| **Video-clip content:Item** | 154608 | 12884.0 | 12 | 453.84 | 41.1526 | < 2.2e-16 *** |
| **Video-clip content:(Stomach) Pressure** | 918 | 229.4 | 4 | 477.11 | 0.7327 | 0.56993 |
| **Video-clip content:(Stomach) pH** | 2335 | 583.8 | 4 | 471.88 | 1.8646 | 0.11549 |
| **Video-clip content:(Stomach) T** | 1905 | 476.3 | 4 | 459.68 | 1.5215 | 0.19478 |
| **Item:(Stomach) Pressure** | 1262 | 420.6 | 3 | 453.84 | 1.3433 | 0.25969 |
| **Item:(Stomach) pH** | 7715 | 2571.7 | 3 | 453.84 | 8.2142 | 2.470e-05 *** |
| **Item:(Stomach) T** | 1216 | 405.3 | 3 | 453.84 | 1.2946 | 0.27568 |
| **Video-clip content:item:(Stomach) Pressure** | 4093 | 341.1 | 12 | 453.84 | 1.0895 | 0.36686 |
| **Video-clip content:item:(Stomach) pH** | 7433 | 619.4 | 12 | 453.84 | 1.9784 | 0.02456 * |
| **Video-clip content:item:(Stomach) T** | 2293 | 191.0 | 12 | 453.84 | 0.6102 | 0.83407 |

**S1. B Results of the Bonferroni corrected post hoc tests** performed, via *emtrends* function, following the significant triple interaction Video-clip content:item:(Stomach) pH, obtained by running the more conservative version of Model 1.

| Video-clip content | Item | pH | SE | df | T ratio | P |
|---|---|---|---|---|---|---|
| **Disgust** | **Disgust** | -15.64 | 5.75 | 159.00 | -2.72 | 0.01 ** |
| **Fearful** | **Disgust** | 6.02 | 3.35 | 152.00 | 1.80 | 0.07 |
| **Happy** | **Disgust** | -2.18 | 2.87 | 235.00 | -0.76 | 0.45 |
| **Neutral** | **Disgust** | -1.68 | 5.17 | 209.00 | -0.32 | 0.75 |
| **Sad** | **Disgust** | 2.98 | 4.38 | 191.00 | 0.68 | 0.50 |
| **Disgust** | **Happy** | 21.12 | 5.75 | 159.00 | 3.67 | <0.001*** |
| **Fearful** | **Happy** | 7.56 | 3.35 | 152.00 | 2.26 | 0.03* |
| **Happy** | **Happy** | 5.94 | 2.87 | 235.00 | 2.08 | 0.04* |
| **Neutral** | **Happy** | 3.55 | 5.17 | 209.00 | 0.69 | 0.49 |
| **Sad** | **Happy** | 6.04 | 4.38 | 191.00 | 1.38 | 0.17 |
| **Disgust** | **Sad** | -2.34 | 5.75 | 159.00 | -0.41 | 0.68 |
| **Fearful** | **Sad** | 1.81 | 3.35 | 152.00 | 0.54 | 0.59 |
| **Happy** | **Sad** | 0.02 | 2.87 | 235.00 | 0.01 | 0.99 |
| **Neutral** | **Sad** | -6.33 | 5.17 | 209.00 | -1.22 | 0.22 |
| **Sad** | **Sad** | 4.68 | 4.38 | 191.00 | 1.07 | 0.29 |
| **Disgust** | **Fear** | -10.90 | 5.75 | 159.00 | -1.90 | 0.06 |
| **Fearful** | **Fear** | 1.02 | 3.35 | 152.00 | 0.31 | 0.76 |
| **Happy** | **Fear** | -0.40 | 2.87 | 235.00 | -0.14 | 0.89 |
| **Neutral** | **Fear** | -0.98 | 5.17 | 209.00 | -0.19 | 0.85 |
| **Sad** | **Fear** | 1.14 | 4.38 | 191.00 | 0.26 | 0.80 |

**Table S2 Results of Model 4 (EGG peak frequency, session 1).** Type III analysis of variance table with Satterthwaite's method.

| | Sum Sq | Mean Sq | NumDF | DenDF | F value | P |
|---|---|---|---|---|---|---|
| **Video-clip content** | 13617 | 3404.3 | 4 | 418.86 | 10.1660 | 7.302e-08 *** |
| **Item** | 8180 | 2726.6 | 3 | 405.51 | 8.1422 | 2.819e-05 *** |
| **Egg peak frequency** | 466 | 465.9 | 1 | 402.48 | 1.3913 | 0.2389 |
| **Video-clip content:item** | 142599 | 11883.3 | 12 | 405.51 | 35.4858 | < 2.2e-16 *** |
| **Video-clip content:egg peak frequency** | 823 | 205.7 | 4 | 425.15 | 0.6142 | 0.6526 |
| **Item:egg peak frequency** | 522 | 173.8 | 3 | 405.51 | 0.5191 | 0.6693 |
| **Video-clip content:item:egg peak frequency** | 2282 | 190.2 | 12 | 405.51 | 0.5679 | 0.8680 |

**Table S3 Results of Model 2 (pill in the small bowel, session 2).** Type III analysis of variance table with Satterthwaite's method.

| | Sum Sq | Mean Sq | NumDF | DenDF | F value | P |
|---|---|---|---|---|---|---|
| **Video-clip content** | 13778 | 3444.6 | 4 | 478.81 | 12.9585 | 5.013e-10 *** |
| **Item** | 3314 | 1104.6 | 3 | 469.26 | 4.1554 | 0.006374 ** |
| **(Small bowel) Pressure** | 78 | 77.7 | 1 | 140.02 | 0.2923 | 0.589630 |
| **(Small bowel) pH** | 805 | 805.1 | 1 | 69.10 | 3.0289 | 0.086244 . |
| **(Small bowel) T** | 43 | 42.7 | 1 | 30.06 | 0.1607 | 0.691327 |
| **Video-clip content:Item** | 208270 | 17355.8 | 12 | 469.26 | 65.2925 | < 2.2e-16 *** |
| **Video-clip content:(Small bowel) Pressure** | 1003 | 250.7 | 4 | 486.53 | 0.9431 | 0.438650 |
| **Video-clip content:(Small bowel) pH** | 778 | 194.4 | 4 | 477.46 | 0.7314 | 0.570808 |
| **Video-clip content:(Small bowel) T** | 181 | 45.4 | 4 | 471.94 | 0.1707 | 0.953324 |
| **Item:(Small bowel) Pressure** | 1160 | 386.7 | 3 | 469.26 | 1.4546 | 0.226235 |
| **Item:(Small bowel) pH** | 610 | 203.5 | 3 | 469.26 | 0.7655 | 0.513789 |
| **Item:(Small bowel) T** | 196 | 65.2 | 3 | 469.26 | 0.2452 | 0.864744 |
| **Video-clip content:item:(Small bowel) Pressure** | 4598 | 383.2 | 12 | 469.26 | 1.4416 | 0.143390 |
| **Video-clip content:item:(Small bowel) pH** | 1930 | 160.8 | 12 | 469.26 | 0.6050 | 0.838509 |
| **Video-clip content:item:(Small bowel) T** | 5273 | 439.4 | 12 | 469.26 | 1.6530 | 0.074340 . |

**TableS4 Results of Model 3 (pill in the large bowel, session 3).** Type III analysis of variance table with Satterthwaite's method.

| | Sum Sq | Mean Sq | NumDF | DenDF | F value | P |
|---|---|---|---|---|---|---|
| **Video-clip content** | 14179 | 3544.7 | 4 | 473.75 | 11.3579 | 8.251e-09 *** |
| **Item** | 4319 | 1439.7 | 3 | 468.86 | 4.6129 | 0.003418 ** |
| **(Large bowel bowel) Pressure** | 3 | 2.8 | 1 | 269.21 | 0.0089 | 0.924857 |
| **(Large bowel bowel) pH** | 0 | 0.0 | 1 | 58.50 | 0.0000 | 0.994501 |
| **(Large bowel bowel) T** | 33 | 33.1 | 1 | 31.27 | 0.1061 | 0.746856 |
| **Video-clip content:Item** | 159021 | 13251.8 | 12 | 468.86 | 42.4611 | < 2.2e-16 *** |
| **Video-clip content:(Large bowel bowel) Pressure** | 1165 | 291.2 | 4 | 485.30 | 0.9331 | 0.444414 |
| **Video-clip content:(Large bowel bowel) pH** | 973 | 243.4 | 4 | 474.95 | 0.7798 | 0.538680 |
| **Video-clip content:(Large bowel bowel) T** | 720 | 180.1 | 4 | 472.93 | 0.5770 | 0.679421 |
| **Item:(Large bowel bowel) Pressure** | 343 | 114.2 | 3 | 468.86 | 0.3659 | 0.777669 |
| **Item:(Large bowel bowel) pH** | 755 | 251.8 | 3 | 468.84 | 0.8069 | 0.490491 |
| **Item:(Large bowel bowel) T** | 938 | 312.6 | 3 | 468.86 | 1.0017 | 0.391836 |
| **Video-clip content:item:(Large bowel bowel) Pressure** | 1496 | 124.7 | 12 | 468.86 | 0.3994 | 0.963735 |
| **Video-clip content:item:(Large bowel bowel) pH** | 2288 | 190.7 | 12 | 468.84 | 0.6109 | 0.833531 |
| **Video-clip content:item:(Large bowel bowel) T** | 2089 | 174.1 | 12 | 468.86 | 0.5578 | 0.875755 |

**Table S5. Emotional video-clips content.** List of emotional video-clips used in the emotional induction task with a brief description in the second column, adapted by Tettamanti et al. 2012.

| Category_Number of the Video-clips | Description of the Video-clips |
|---|---|
| Disgust_1 | A monstrous creature vomits a corrosive fluid on a dead man's face and dissolves it |
| Disgust_2 | A severed human head is covered with insects |
| Disgust_3 | A man is covered with faecal matter |
| Disgust_4 | Some dishes and a wall are covered with insects |
| Disgust_5 | A man takes a worm off an animal's wound and holds it |
| Disgust_6 | A monstrous creature vomits a corrosive fluid on a dead man's foot |
| Disgust_7 | A monstrous creature vomits a corrosive fluid on a dead man's hand and dissolves it |
| Disgust_8 | A man plunges his hand in a toilet, retracts it covered with faecal matter and almost vomits |
| Disgust_9 | A man removes part of his own skin |
| Disgust_10 | A man vomits in a bowl, a man watches it and another man smells it |
| Disgust_11 | A man takes off his fingernail |
| Disgust_12 | A man eats snails |
| Disgust_13 | Close up on a fly which approaches a severed human hand covered with insects |
| Disgust_14 | Close up on insects |
| Disgust_15 | A man smells a bowl of vomit and drinks it |
| Disgust_16 | A girl vomits on a man's face |
| Disgust_17 | A man squeezes a purulent dot on his hand |
| Disgust_18 | A man is covered with a substance and tries to remove it from his skin |
| Disgust_19 | Some surgeons perform a surgical operation on a bloodied inhuman creature |
| Disgust_20 | A monstrously shaped egg cracks open and viscous substances spill out |
| Disgust_21 | A bloodied girl whose upper lip is missing bites a man's neck |
| Disgust_22 | A purulent inhuman organ is being dissected |
| Disgust_23 | A woman vomits on a man's face |
| Disgust_24 | A monstrous creature is dissected |
| Fear_1 | A girl is approached by a scary creature |
| Fear_2 | A bear roars against a man |
| Fear_3 | A man attacks a woman |
| Fear_4 | A woman is approached by a monster |
| Fear_5 | Several men are falling from a mountain |
| Fear_6 | A woman is held underwater by a scary creature |
| Fear_7 | A woman tries to escape and she is stabbed in her hand |
| Fear_8 | A woman is attacked by a man |
| Fear_9 | A woman is attacked by a scary creature |
| Fear_10 | A man is attacked by a shark |
| Fear_11 | An aeroplane is disrupted during flight |
| Fear_12 | A woman is attacked by an alien |
| Fear_13 | A man is attacked by a shark |
| Fear_14 | A man is tied to a chair with his mouth duct taped |
| Fear_15 | Two climbers hang in the void |
| Fear_16 | A woman is attacked by a man |
| Fear_17 | A woman falls from a mountain |
| Fear_18 | A woman falls from a building |
| Fear_19 | A man is attacked by a man |
| Fear_20 | A man at risk of being crashed |
| Fear_21 | Two man hang on a building ledge |
| Fear_22 | A woman is approached by a monster |
| Fear_23 | A woman is attacked by a man |
| Fear_24 | A man is trapped underwater |
| Happy_1 | Several persons rejoice at a soccer match |
| Happy_2 | Several boys and a man rejoice after scoring a goal in a soccer match |
| Happy_3 | A woman smiles after seeing an old lady beating a young man with her purse |

| | |
|---|---|
| Happy_4 | A child smiles while looking at two persons smiling and hugging each other |
| Happy_5 | A woman smiles while looking at a man playing with a child |
| Happy_6 | A woman and a man kiss each other, women claps their hands |
| Happy_7 | A woman and a man run on a shore playing with each other |
| Happy_8 | A man rejoices after scoring a goal in a soccer match |
| Happy_9 | A woman and a man kiss each other |
| Happy_10 | A woman and a man kiss each other |
| Happy_11 | A woman and a man play together |
| Happy_12 | A man and a woman laugh together |
| Happy_13 | A man hugs a woman |
| Happy_14 | A man and a woman smile while looking at a dog lactating her puppies |
| Happy_15 | Several persons dance and a girl laughs |
| Happy_16 | A woman and a man laugh together |
| Happy_17 | A girl opens a present, a woman kisses her |
| Happy_18 | A man scores a goal in a soccer match |
| Happy_19 | A man hugs his sons |
| Happy_20 | A woman and a man dance and smile at each other |
| Happy_21 | Several persons rejoice at a soccer match |
| Happy_22 | Players celebrating for scoring a goal |
| Happy_23 | Several persons rejoice at a soccer match |
| Happy_24 | Several persons rejoice at a rugby match |
| Sad_1 | A woman cries and holds the hand of a man in a hospital bed |
| Sad_2 | A man cries while holding a dead woman |
| Sad_3 | Two persons are separated as the train leaves |
| Sad_4 | A sick child is in a hospital bed |
| Sad_5 | A man cries while holding a dead woman |
| Sad_6 | A woman kisses the forehead of a dead boy |
| Sad_7 | A woman cries on the phone |
| Sad_8 | A woman cries while holding a dead man |
| Sad_9 | Dead bodies are moved by the waves on the shore |
| Sad_10 | A man cries while holding the bloodied body of a man |
| Sad_11 | A woman cries and hugs a man |
| Sad_12 | A sick woman is in bed, a man holds her hand |
| Sad_13 | A man crying while looking at a dead man |
| Sad_14 | A woman cries, two women hug her |
| Sad_15 | A man holds a dead man |
| Sad_16 | Several dead bodies lay upon the ground |
| Sad_17 | Several persons are in a hospital bed |
| Sad_18 | Dead bodies lay on a shore |
| Sad_19 | Several persons watch three coffins on the ground |
| Sad_20 | A woman and a man hold each other |
| Sad_21 | Three persons cry while holding each other |
| Sad_22 | A man looks upon a dead body |
| Sad_23 | A woman closes herself in a room and cries |
| Sad_24 | A woman lays a flower on a coffin at a funeral, another woman watches |
| Neutral_1 | Two men, one speaking and the other one listening |
| Neutral_2 | A woman opening the window of the dog kennel in an old mansion |
| Neutral_3 | A woman meets a man at the train station |
| Neutral_4 | A woman takes the computer from a bag and puts it on a table |
| Neutral_5 | Several persons observe drawings and sculptures |
| Neutral_6 | Several persons talk and walk in an office |
| Neutral_7 | A woman walks in a large building, several other persons in the background |
| Neutral_8 | Two waiters greet several persons entering a room |
| Neutral_9 | A man closes the door and walks away |
| Neutral_10 | A man works in a building site, several other men working in the background |
| Neutral_11 | A woman in a boat sits and opens a book |
| Neutral_12 | Several persons walking in a corridor, a woman enters an office |
| Neutral_13 | Men carving and moving large cinematographic props |
| Neutral_14 | A woman and a man speak while cleaning the kitchen |
| Neutral_15 | A woman walks down the road, some children on bikes wave at her |

| | |
|---|---|
| **Neutral_16** | A women's meeting, one speaks while the others nod |
| **Neutral_17** | A woman hoovers a room |
| **Neutral_18** | A man walks in the countryside while reading |
| **Neutral_19** | A man writes while sitting on the outside of a ferry |
| **Neutral_20** | Different scenes from a food market |
| **Neutral_21** | A woman types at the computer and lifts the receiver |
| **Neutral_22** | A man and two young men talk by the phone |
| **Neutral_23** | A man drawing |
| **Neutral_24** | A man cleans the leaves off the yard |
